# Why does Δ^9^-Tetrahydrocannabinol act as a partial agonist of cannabinoid receptors?

**DOI:** 10.1101/2021.04.29.441987

**Authors:** Soumajit Dutta, Balaji Selvam, Aditi Das, Diwakar Shukla

**Affiliations:** Department of Chemical and Biomolecular Engineering, University of Illinois at Urbana-Champaign, Urbana, IL, 61801; Department of Comparative Biosciences,University of Illinois at Urbana-Champaign, Urbana, IL, 61802; Department of Bioengineering,University of Illinois at Urbana-Champaign, Urbana, IL, 61801; Department of Biochemistry,University of Illinois at Urbana-Champaign, Urbana, IL, 61801; Center for Biophysics and Quantitative Biology, University of Illinois at Urbana-Champaign, Urbana, IL, 61801; Cancer Center at Illinois, University of Illinois at Urbana-Champaign, Urbana, IL, 61801; National Center for Supercomputing Applications, University of Illinois, Urbana, IL, 61801; Beckman Institute for Advanced Science and Technology, University of Illinois at Urbana-Champaign, Urbana, IL, 61801; NIH Center for Macromolecular Modeling and Bioinformatics, University of Illinois at Urbana-Champaign, Urbana, IL, 61801

## Abstract

Cannabinoid receptor 1 (CB_1_) is a therapeutically relevant drug target for controlling pain, obesity, and other central nervous system disorders. However, full agonists and antagonists of CB_1_ have been reported to cause serious side effects in patients. Therefore, partial agonists have emerged as a viable alternative to full agonists and antagonists as they avoid overstimulation and side effects. One of the key bottlenecks in the design of partial agonists is the lack of understanding of the molecular mechanism of partial agonism. In this study, we examine two mechanistic hypotheses for the origin of partial agonism in cannabinoid receptors and explain the mechanistic basis of partial agonism exhibited by Δ^9^-Tetrahydrocannabinol (THC). In particular, we inspect whether partial agonism emerges from the ability of THC to bind in both agonist and antagonist binding pose or from its ability to only partially activate the receptor. Extensive molecular dynamics simulations and the Markov state model capture the THC binding in both antagonist, and agonist binding poses in CB_1_ receptor. Furthermore, we observe that binding of THC in the agonist binding pose leads to rotation of toggle switch residues and causes partial outward movement of intracellular transmembrane helix 6 (TM6). Our simulations also suggest that the alkyl side chain of THC plays a crucial role in determining partial agonism by stabilizing the ligand in the agonist and antagonist-like poses within the pocket. This study provides us fundamental insights into the mechanistic origin of the partial agonism of THC.

## Introduction

Cannabinoid receptor 1 belongs to the family of Class A G-Protein Coupled Receptors (GPCRs) (*1, 2*), which modulates diverse cellular signaling processes via intracellular G-proteins (*3*) and *β*-arrestins (*4*). CB_1_ receptors were first discovered in the last decade of the twentieth century as a target of plant cannabinoid molecules (*5*). Due to its ubiquitous presence in physiological processes, CB_1_ is an important drug target for the potential treatment of a variety of diseases. In the last thirty years, several synthetic molecules have been designed to target the CB_1_ for the treatment of pain, obesity, and inflammation (*6–8*). CB_1_ receptor agonists such as MDMB-Fubinaca (*3*) and inverse agonists such as Rimonabant (*9, 10*) have been shown to modulate the receptor activity significantly. However, these designed agonists and antagonists of CB_1_ also exhibit dangerous side effects. For instance, Fubinaca, also known as “zombie drug”, caused the hospitalization of thousands of patients in New York (*11*). Rimonabant had to be withdrawn from the market due to its psychotic side effects like depression and anxiety (*12*). While these designed agonists and antagonists have failed to meet the drug safety guidelines, alternate approaches (e.g., allosteric modulator, partial agonist) can be explored for designing therapeutics. We recently studied the binding of a negative allosteric modulator (NAM), sodium ion (Na^+^), to cannabinoid receptors (CBs) using molecular dynamics simulation (*13*). Simulation revealed important differences in binding site and pathway between CB_1_ and CB_2_, which can be exploited to design a selective NAM drug. Similarly, a partial agonist Dronabinol has been approved by Food and Drug Administration (FDA) as an appetite stimulant drug for AIDS patients and an antiemetic drug for chemotherapy (*14*). Dronabinol is a synthetic form of THC (Figure 1), the main psychoactive compound in marijuana which binds to the CB_1_ as a partial agonist and affects the endocannabinoid signaling pathway. THC has been shown to demonstrate positive effects for treating Huntington’s disease, Parkinson’s disease, Alzheimer’s disease (*15*). Although THC has the potential to become a valuable drug for several diseases, this drug is still banned by the FDA due to its side effects. Therefore, molecular level understanding of THC and the mechanism by which it partially activates the cannabinoid receptors will inform the design of potential partial agonist drugs targeting CB_1_ receptor. Structural studies of CB_1_ have revealed that toggle switch residue (TRP356^6.48^ and PHE200^3.36^) movement by the agonist is crucial for the activation of CB_1_ (Figure 2B)(*3, 16–21*). Kumar et al.(*3*) proposed that due to the smaller size of THC as compared to the full agonists, there is less interaction between toggle switch residues and ligand. Furthermore, they proposed that the THC binding position with downward facing alkyl chain can shift the toggle switch residues and activate the receptor. However, the crystal structures and docking studies only provide static interactions of a ligand in the binding pose. These studies do not reveal the mechanism of ligand conformational switching inside the binding pocket. This proposed hypothesis has not been rigorously examined from either the experimental or computational approaches. Therefore, it is difficult to obtain structural understanding of the origin of partial agonism exhibited by THC. A recent study employing Metadynamics simulations showed that a partial agonist GAT228 binds in multiple positions inside the ligand binding pocket of CB_1_ (*22*) due to the large size of the pocket as compared to the ligand volume (*18*). This observation is also consistent with the distinct poses of agonist and antagonist in the binding pocket (Figure 2A). Therefore, it is likely that partial agonist THC might be stabilized in both the agonist and antagonist-like pose inside the binding pocket and only the subset of THC molecules bound to CB_1_ in the agonist-like pose activate the receptor. This phenomenon would decrease the maximum response of the secondary messenger. The first hypothesis is shown as equations 1, 2, 3 where R, RA**, RPA**,[RPA] represent the receptor in apo form, agonist bound active state, partial agonist bound active and inactive state, respectively. In RPA** and [RPA], partial agonist binds in agonist and antagonist bound poses, respectively. Agonist and partial agonist are represented as A and PA.

**Figure 1:**
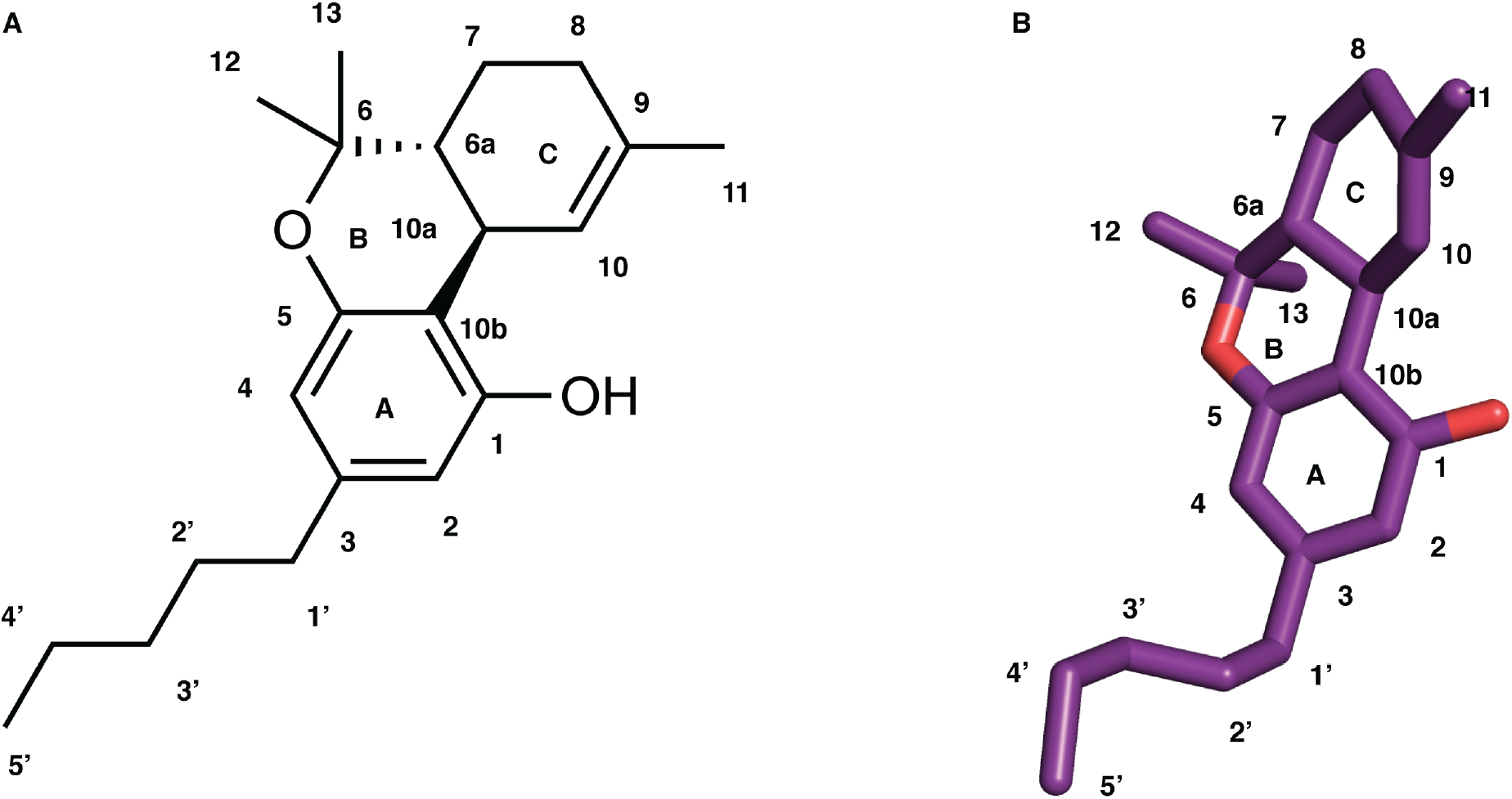
2D and 3D representation of THC structure. Numbering of carbon atoms and rings are mentioned in the figure.

**Figure 2:**
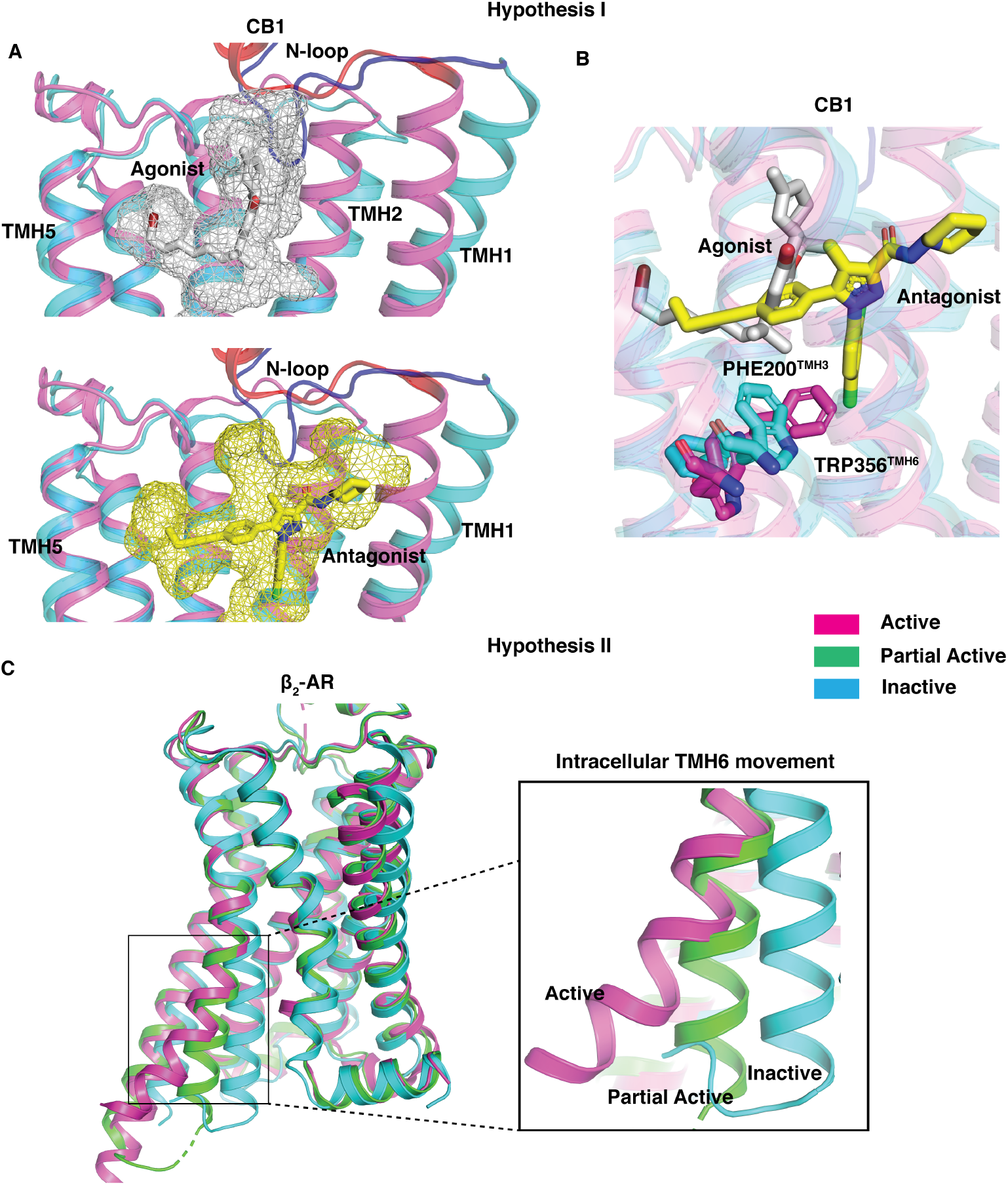
Pictorial representation of both the hypotheses. (A), (B) are representing hypothesis 1. Superposition of active (PDB ID: 5XRA, color: pink) and inactive (PDB ID: 5TGZ, color: cyan) structures of CB_1_ is shown in all panels of (A). In the top and bottom panel of (A), antagonist (AM6538, color: yellow) and agonist (AM11542, color: silver) are shown as stick and mesh representation. N-loop is colored differently (Active: red, Inactive: Blue). TM6 and TM7 are not shown in the cartoon representation for better visualization of ligand poses. In (B) toggle switch residues (TRP356^6.48^ and PHE200^3.36^) are shown as sticks. Superposition of active (PDB ID: 3SN6, color: pink), partially active (PDB ID: 6CSY (*23*), color: green), and inactive (PDB ID: 2RH1, color: cyan) structures of *β*_2_-AR is shown in (C). Intrcaellular TM6 movement is highlighted in a separated box.

### Hypothesis I

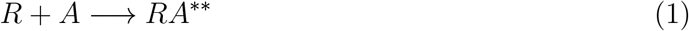

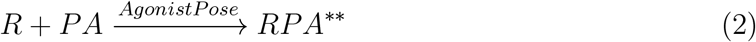

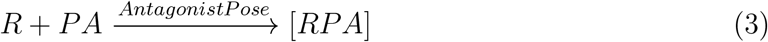

Canonical class A GPCR activation is characterized by intracellular TM6 movement, which facilitates the G-protein binding. A partial agonist, Salmeterol, binds in the orthosteric pocket in the *β*_2_-Adrenergic receptor and causes partial movement of TM6 compared to the full movement by an agonist (Figure 2C) (*23*). Therefore, we hypothesize that partial activation may happen if the partial agonist stabilizes the receptor in different conformation than the active structure. THC has smaller side chain compared to agonist AM11542 (*18*). Furthermore, absence of the dimethyl group at the first carbon (C1’) of the alkyl chain decrease the interaction with toggle switch residues. Thus, we propose that THC binding may also cause the partial outward movement of the TM6.

### Hypothesis II

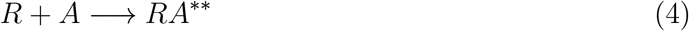

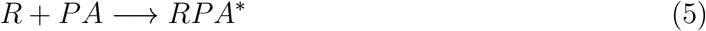

The second proposed hypothesis is explained in equation 4, 5 where RPA* represents partial agonist bound partially active form of the receptor. Other notations are similar to the reactions 1, 2, 3.

We assess the validity of these hypotheses by running extensive simulations of THC binding to CB_1_ (see methods section). Using the Markovian property of Molecular Dynamic (MD) simulations, we build Markov State Model (MSM) using the simulation data. From the eigenvalues and vectors of MSM, we obtain the timescale and the thermodynamics of the ligand binding process. MSM weighted data reveals that THC is stabilized in both antagonist and agonist bound poses in agreement with our first hypothesis. The free energy barrier of ~ 1kcal/mol is estimated for THC to transition between the antagonist to agonist bound pose, which implies that thermal fluctuations at the human body temperature could easily allow the THC to transition between these poses. In the agonist bound pose, THC rotates the important toggle switch residue TRP35 6^6.48^. The new position of the TRP356^6.48^ is different compared to active state of CB_1_. This intermediate position TRP356^6.48^ leads to partial outward movement of TM6 suggesting THC can only partially active the receptor according to out second hypothesis. This mechanistic study explains the reason behind the partial agonism behavior of THC compared to other agonists and will aid future drug development targeting cannabinoid receptors.

## Results

### THC is stabilized in both antagonist and agonist binding poses in the orthosteric pocket of CB_1_

Active and inactive structures of CB_1_ reveal that orthosteric binding site volume undergoes a large change upon activation as compared to other class A GPCRs (*18*). Comparison of active (PDB ID: 5XRA (*17*)) and inactive (PDB ID: 5TGZ (*18*)) structures also reveal that the agonist and antagonist bind in the different regions within the pocket (Figure 2A). Agonist molecule binds in a region close to the TM5, whereas antagonist binds in the extended pocket formed by TM1 and TM2. In the inactive structure, the downward movement of the N-loop towards the binding pocket separates the agonist and antagonist binding regions (Figure 2A). However, partial agonist bound crystal structure of CB_1_ or CB_2_ is not reported in the literature. Therefore, the partial agonist binding position is not well documented. Preliminary docking studies in both inactive and active structures reveal that THC binds to the agonist binding region in a similar conformation as other agonists (*17, 18*). However, docking alone cannot infer the exact binding pose of a ligand which undergoes a dynamic conformational change within the binding pocket. Therefore, we perform ~ 143*μ*s of MD simulations to characterize the THC binding mechanism in CB_1_ starting with inactive structure.

To capture the THC binding process and upward movement of the N-loop, we project the MD simulation data along the two metrics that characterize these motions (Figures 3A, S1A and S1B). MSM weighted free energy landscape plot (Figure 3A) shows the movement of THC molecule towards TM5a s indicated by the distance between THC-C1’ atom (Figure 1) and TYR275^5.39^-C*α* atom on TM5. THC enters CB_1_ binding pocket through the space between the space of TM1, TM2, and N-loop (Figure S2) (*24*). THC further diffuses inside the pocket and is stabilized in antagonist binding pose where distance between THC(C1’) and TYR275^5.39^(C*α*) is 15 to 21 Å(Figure 3B). We observe two stable local energy wells in the antagonist binding pose. The free energy well away from agonist bound pose is named antagonist-like pose 1 and second well is named antagonist-like pose 2. These two minima are separated by the activation barrier of 0.55 ± 0.43 kcal/mol. Superposition of predicted MD structures of THC bound in antagonist-like pose 1 and 2 with inactive structure shows that B and C ring of the tricyclic dibenzopyran group of THC binds in same position as Arm 3 of the antagonist, AM6538 (Figures 3C and 3D) (*17*). The aromatic ring of THC matches with pyrazole ring of the antagonist. However, THC alkyl chain (side chain) orientation varies between the two binding poses. In antagonist-like pose 1, it extends towards the conserved sodium binding site (*13, 25, 26*) similar to the arm 1 of antagonist, whereas, in antagonist-like pose 2, it orients itself in the direction of agonist binding site similar to the arm 2 of the antagonist (Figures 3C and 3D). In both the antagonist-like poses, THC forms stable polar interaction with SER383^7.39^ and hydrophobic interaction with N-loop (PHE102^N–loop^, MET103^N–loop^), TM1(SER123^1.39^, ILE119^1.35^), TM2 (PHE170^2.57^) and TM7 (ALA38 0^7.36^) (Figures 4B, 4C, S3A and S2B). In these poses, N-loop remains inside the pocket and restricts the movement of THC towards the agonist binding region. The upward movement of N-loop allows the movement of THC inside the pocket and stabilizes it in the agonist-like pose (Figure 3E). THC has smaller alkyl side chain than cannabinoid like full agonists of CB_1_, thus, first carbon of THC (C1’) binds closer to to TM5 compared to the agonist, AM11542. In agonist binding pose, THC(OH) forms polar interaction with SER383^7.39^ same as other agonists. Furthermore, THC forms extensive hydrophobic interactions with amino acid residues of TM3 (VAL196^3.32^, LEU193^3.29^), TM5 (TRP279^5.43^), TM7 (PHE379^7.35^), N-loop (MET103^N–loop^), and ECL2 (PHE268^ECL2^) as shown in Figures 4D and S3C.

**Figure 3:**
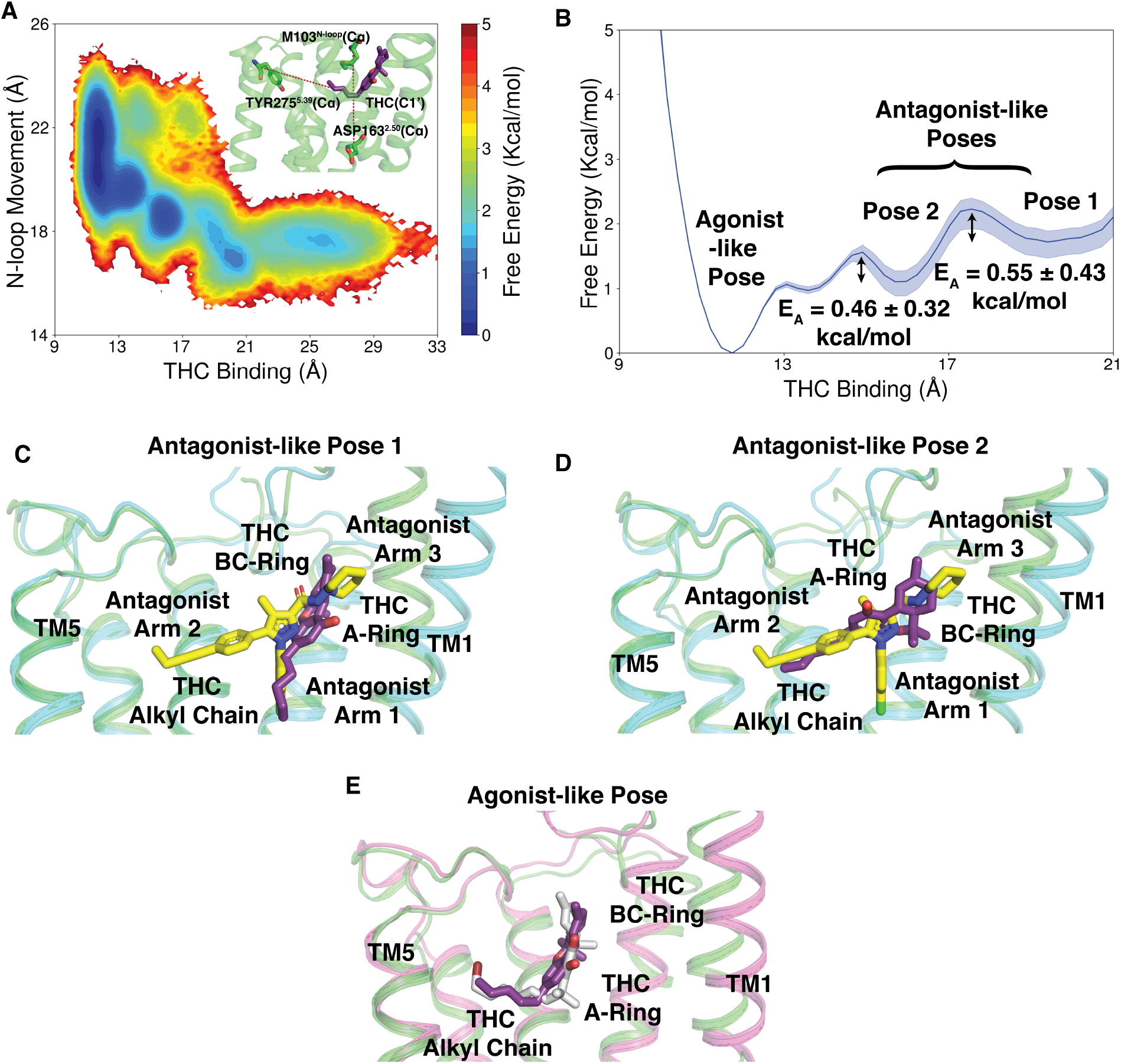
Distinct stabilized poses of THC inside binding pocket of CB_1_. (A) MSM weighted free energy landscape to capture THC binding and N-loop upward motion. THC binding distance is measured between THC-C1’ and TYR275^5.39^-C*α* (TM5) and N-loop upward motion is measured between MET103^N–loop^-C*α* (N-loop) and ASP163^2.50^-C*α* (TM2). (B) One dimensional free energy diagram depicting stabilized binding position of THC and activation barrier between them. (C), (D) Superposition of inactive (PDB ID: 5TGZ (*17*), color:cyan) structure of CB_1_ and MD snapshots from antagonist-like pose 1 (C) and 2 (D). (E) Superposition of active (PDB ID: 5XRA (*18*), color:pick) structure of CB_1_ and MD snapshot from agonist-like pose. MD snapshots are shown in green color. Agonist (AM11542 (*18*)), partial agonist (THC) and antagonist (AM6538 (*17*)) are represented as sticks with silver, violet, and yellow color respectively. TM6 and TM7 are not shown in the cartoon representation for better visualization of ligand poses.

**Figure 4:**
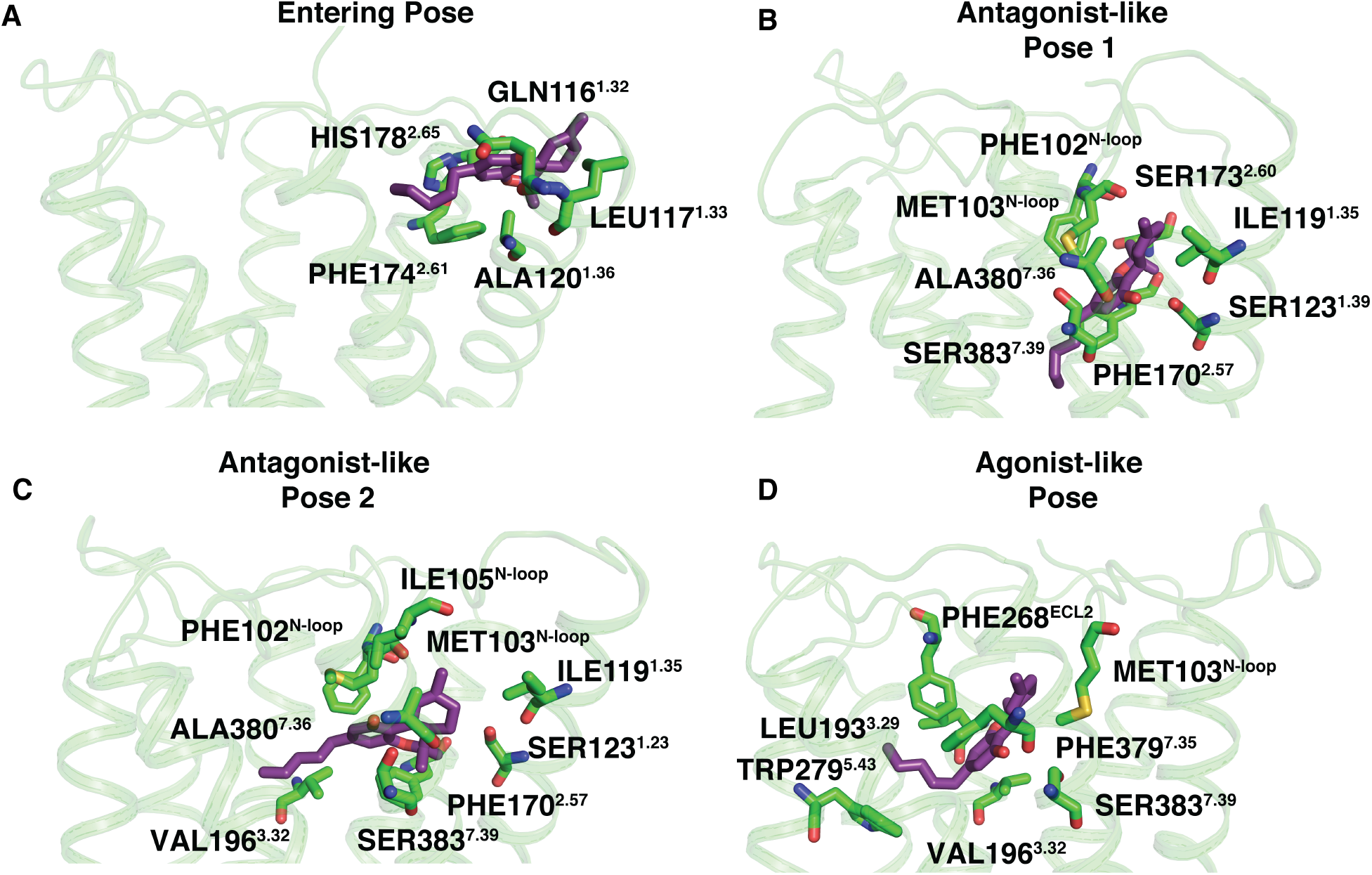
Important interactions between protein residues and THC at different stabilized positions during binding (side view). (A) A representative structure when THC enters the receptor through the space between N-loop, TM1 and TM2. (B), (C) Representative structures from antagonist-like pose 1 and 2, respectively. (D) A representative structure from agonist-like pose. Stable interactions were measured using GetContacts package. Protein structures are shown as cartoon representation (color:green). THC (color:violet) and interactive residues (color:green) are shown as stick.

To capture the timescale of this entire binding process, we run Kinetic monte carlo (kMC) simulations on MD data. kMC utilizes MSM transition probability matrix to find the probable pathway for binding (method section). 150 *μ*s long KMC trajectory reveals that entire binding process from unbound to bound poses takes approximately 100 *μ*s (Figure S4A). From the solution, THC is first stabilized in antagonist bound pose in ~ 50*μ*s. THC occupies antagonist pose I and II for approximately 30 *μ*s and subsequently moves to the agonist binding pose (Figure S4A).

### THC chain orientation plays an important role in partial agonism

Alkyl chain of THC plays an important role in binding and activation of CB_1_. Modification of Alkyl chain leads to change in binding affinity. For example, increasing the chain length of Δ^8^-THC (structural homolog of Δ^9^-THC) from five carbons to eight carbons increases the binding affinity from ~40 nM to ~8nM (*27*). Furthermore, adding a dimethyl group in first carbon of the alkyl chain is hypothesized to increase the interaction with toggle switch TRP356^6.48^ (*18*). To characterize the importance of the alkyl chain of THC, we observe chain dihedral angle (C2-C3-C1’-C2’) (Figure 1) movement during binding. Positive dihedral is crucial to orient the alkyl chain of THC in orthogonal direction of aromatic group as in agonist-like pose (Figure 3E). The free energy landscape of THC alkyl chain dihedral with respect to binding (Figures 5A and S5) reveals that THC binds to CB_1_ in two different chain orientations. If hydrogen atom on C1 faces downward (or positive dihedral angle) during the binding, it moves to agonist binding pocket with maximum free energy barrier of ~ 2 ± 0.4 kcal/mol. This alkyl chain orientation enables THC to pressurize N-loop in upward direction to take orthogonal conformation similar to full agonist. However, if THC enters the receptor with negative dihedral, it is stabilized in the antagonist-like poses. High free energy barrier of ~ 4 ± 0.4 kcal/mol is required for THC to move from macrostate 3’ to 4’ (Figure 5A). In this conformation (macrostate 3’), tricyclic dibenzopyran group of THC is unable shift N-loop upward as the active structure. Hence, alkyl chain of THC shifts the population to macrostate 3 which is more accessible as the free energy barrier is lower than eariler transition. Applying Transition path theory (TPT) on MSM states we can calculate effective timescales of these macrostate transitions. TPT provides mean free passage time (MFPT) for the transition between two macrostates by taking into account all possible pathways through the intermediate states. Calculated MFPT shows that transition between state 3’ to 3 (2.4 ± 0.5*μ*s) is more accessible compared to the 3’ to 4’ (13.2 ± 3.5*μ*s) (Figures 5B and S6). The binding of THC to the agonist-like pose, shows the orientation of alkyl chain may flip between macrostate 4 to 4’ as they are energetically favorable. However, THC has more tendency to stay in the orthogonal conformation (macrostate 4) compared to macrostate 4’. The conditional probability of THC to be in orthogonal conformation in the agonist-like pose is 0.9 ± 0.0. Our results show that THC is stabilized in agonist and antagonist-like poses that supports our first hypothesis.

**Figure 5:**
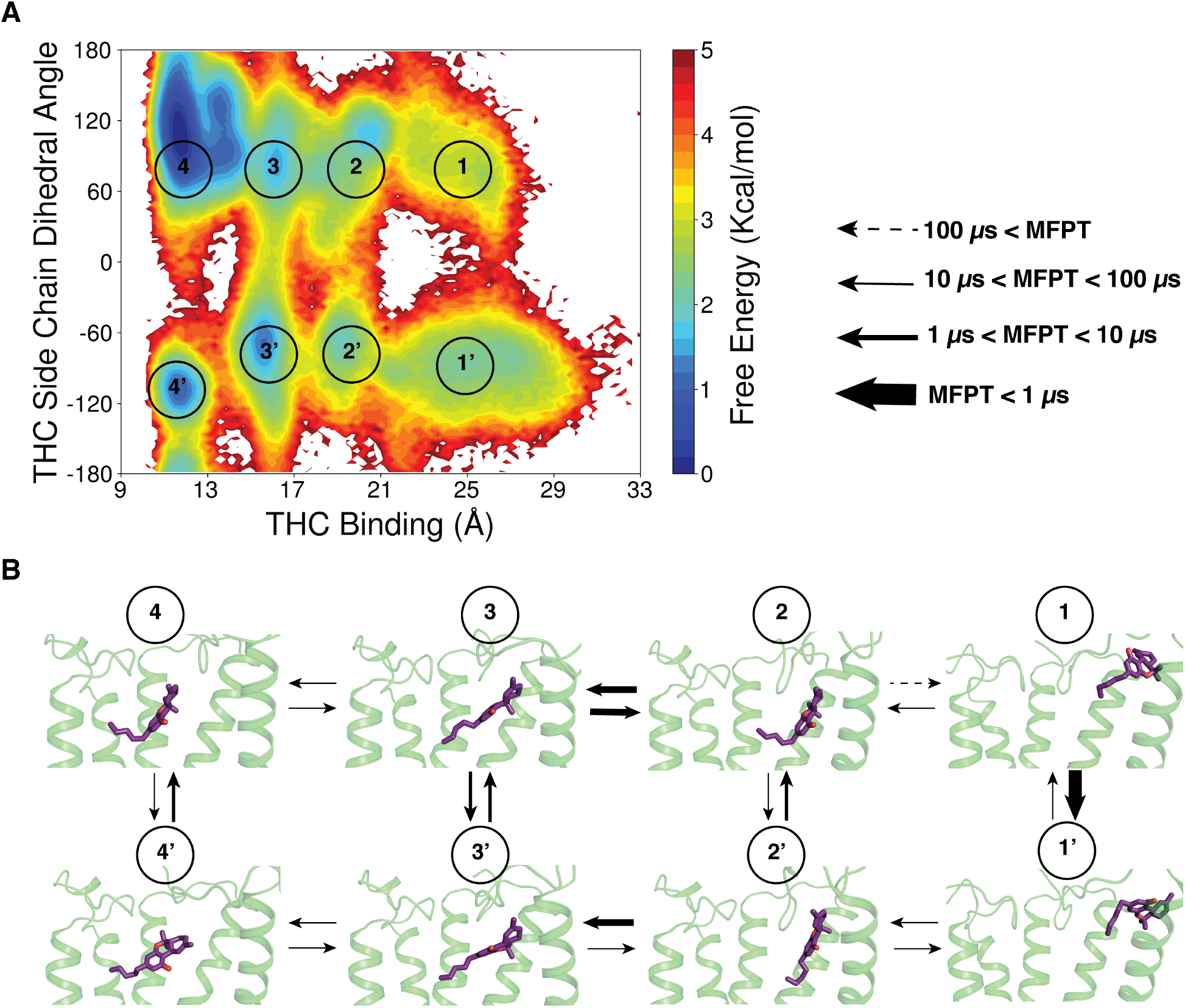
THC alkyl chain movement during binding to CB_1_. (A) MSM weighted free energy landscape to capture THC binding and THC side chain dihedral. THC binding distance is measured between THC-C1’ and TYR275-C*α* (TM5) and THC sidechain dihedral is measured between C2,C3,C1’,C2’. Manually defined macrostate regions are numbered in the figure. (B) Mean free passage time (MFPT) of transitions between the 8 macrostates are shown. Each macrostate is represented by MD snapshot from the region. Different range of MFPT are shown with distinguished arrow thickness. Protein structures are shown as cartoon and THC molecules are shown as stick. TM6 and TM7 are not shown in the cartoon representation for better visualization of ligand poses.

### THC rotates toggle switch TRP356^6.48^ in the agonist binding pocket

Although, we establish the fact that THC can be stabilized in different position of the orthosteric binding pocket, it is not clear how THC activates the receptor. Crystal structures of the CB_1_ receptor in active and inactive position depicts that toggle switch residues play an important role in the activation of the receptor. Agonist molecule triggers the movement of toggle switch residue TRP356^6.48^ movement of the receptor towards the TM5 which consequently leads to outward movement of TM6 (*18*). It was hypothesized that THC behaves as partial agonist due to the lack of interaction with toggle switch residues (*3*). However, we observe that binding of THC leads to the rotation of TRP356^6.48^ (Figures 6A and S7). The dihedral angle of TRP356^6.48^ shifts from inactive conformation (*χ*_2_ dihedral angle between 60° to 120°) to new intermediate active conformation (−30° to 30°) as well as relatively less stable state (−120° to −60°) (Figures 6A and 6B). We referred energetically favorably accessible states as partially active state 1 and state 2, respectively. The rotation of TRP356^6.48^ leads to breakage of aromatic interactions with PHE200^3.36^ and it moves towards TM2 similar to active structure (Figure 6B). Comparison of representative structures from partially active state 1 with inactive CB_2_ shows that toggle switch TRP356^6.48^/TRP258^6.48^ has similar rotamaric conformation as CB_2_ inactive structure (Figure S8). An inverse agonist of CB_2_, MRI2687, which was predicted to retain the toggle switch residue to similar conformation (*28*), acts as a partial agonist for CB_1_. This in turn supports our prediction that THC stabilizes the TRP^6.48^ in intermediate state to partially activate the receptor.

**Figure 6:**
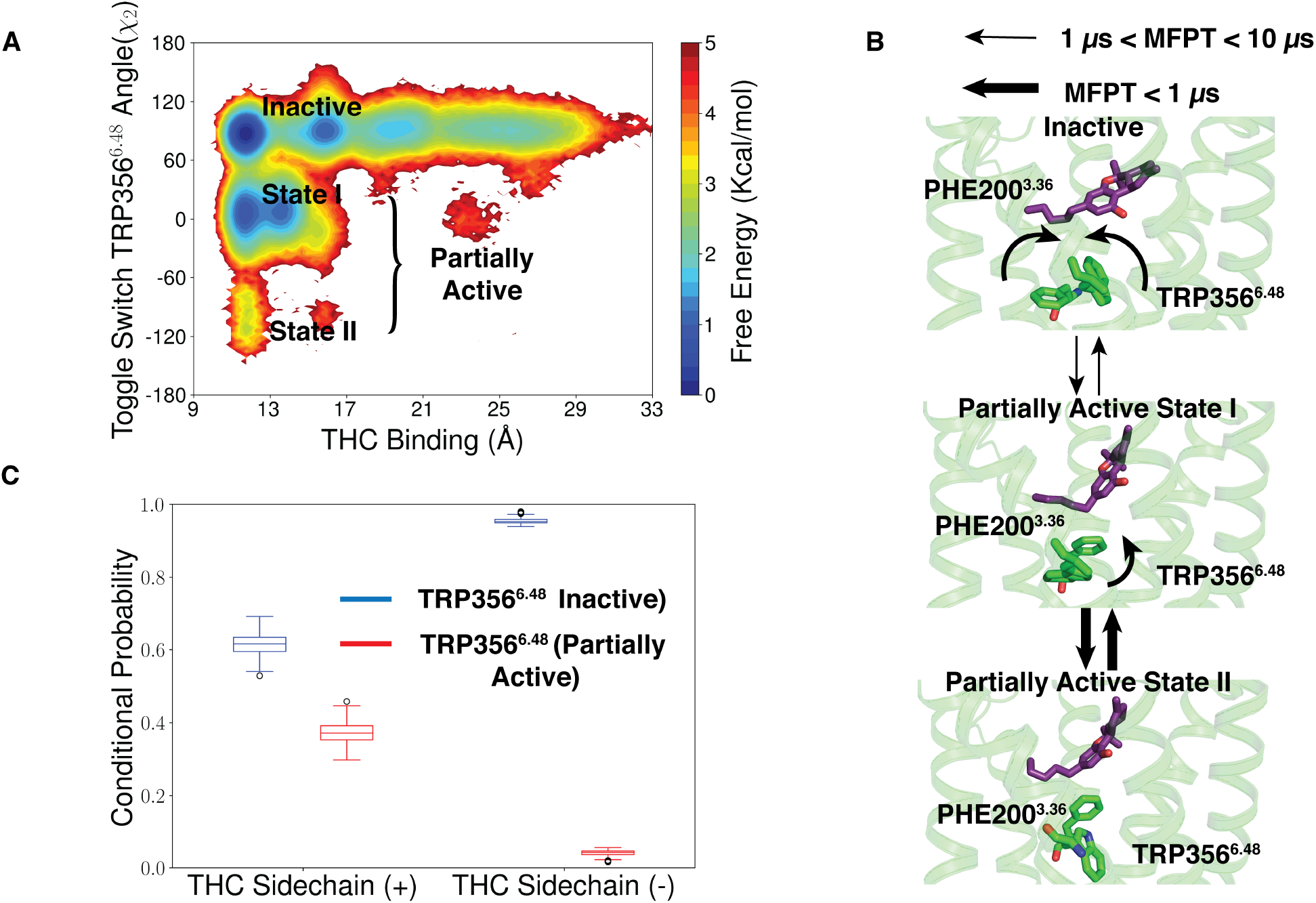
TRP356^6.48^ rotation due to THC binding in Agonist-like pose (A) MSM weighted free energy landscape to capture THC binding and toggle switch TRP356^6.48^ *χ*_2_ angle. THC binding distance is measured between THC-C1’ and TYR275^5.39^-C*α* (TM5). Inactive and partially active (state 1 and 2) macrostates are marked when the THC is bound to the agonist bound pocket. (B) Mean free passage time (MFPT) of transitions between the inactive to partially active conformations of toggle switch residues are shown. Direction of the conformational change in TRP356^6.48^ and PHE200^3.36^ are shown via arrow. Different range of MFPT are shown with distinguished arrow thickness. Protein structures (color: green) are shown as cartoon. THC molecules (color: violet) and toggle switch residues (color: green) are shown as stick. (C) Box plot to show the probability of TRP356^6.48^ rotation with positive or negative THC dihedral in agonist-like pose. Blue and red boxes show the conditional probabilities when TRP356^6.48^ is in inactive and partially active pose. Data distribution in box plot is generated with 200 rounds of bootstrap sampling with 80% of total number of trajectories (Method section).

Although, THC can rotate toggle switch TRP356^6.48^ for subset of structures in agonist binding position, we noticed a favorable free energy minima around inactive pose of TRP356^6.48^. In these minima, THC is unable to shift the toggle switch movement. To explain the reason for the two different orientation of TRP35 6^6.48^, we calculated the probability TRP356^6.48^ rotation with respect to the THC alkyl chain dihedral in agonist-like pose (Figure 6C). Our calculations show that with negative alkyl chain dihedral, THC can move TRP356^6.48^ for only 4.0 ± 0.8% structures. With negative dihedral, the first carbon of the alkyl chain (C1’) binds far from the toggle switch and cannot induce conformational change (Figure 5B). However, binding with a positive dihedral leads to rotation of TRP35 6^6.48^ for 95.5 ± 0.8% of the structures due to more interactions.

The changes in the toggle switch residue leads to the outward movement of TM6 as shown in the Figures 7A and S9. When the toggle switch remains in the inactive position, intracellular TM6 can move easily 2-3 Åeither side inactive structure (low free energy barrier). For this case, TM3-TM6 distance is similar or less than the inactive structure with a probability of 0.74 ± 0.0 (Figures 7B). However, TRP356^6.48^ rotation creates a torsion in TM6 and stabilizes the intracellular part of TM6 of the receptor in between inactive and active-like conformation with probability of 0.6 ± 0.0 (Figures 7B). Comparison of representative structure from partially active state minima with *β*_2_-AR inactive structure shows that intracellular TM5, TM6, and TM7 matches well (Figure S10). Therefore, THC in partially activated state forms favorable interaction with TRP356^6.48^ and stabilize in intermediate conformation compared to the inactive structure. These results support our second hypothesis that THC can only activate the receptor partially.

**Figure 7:**
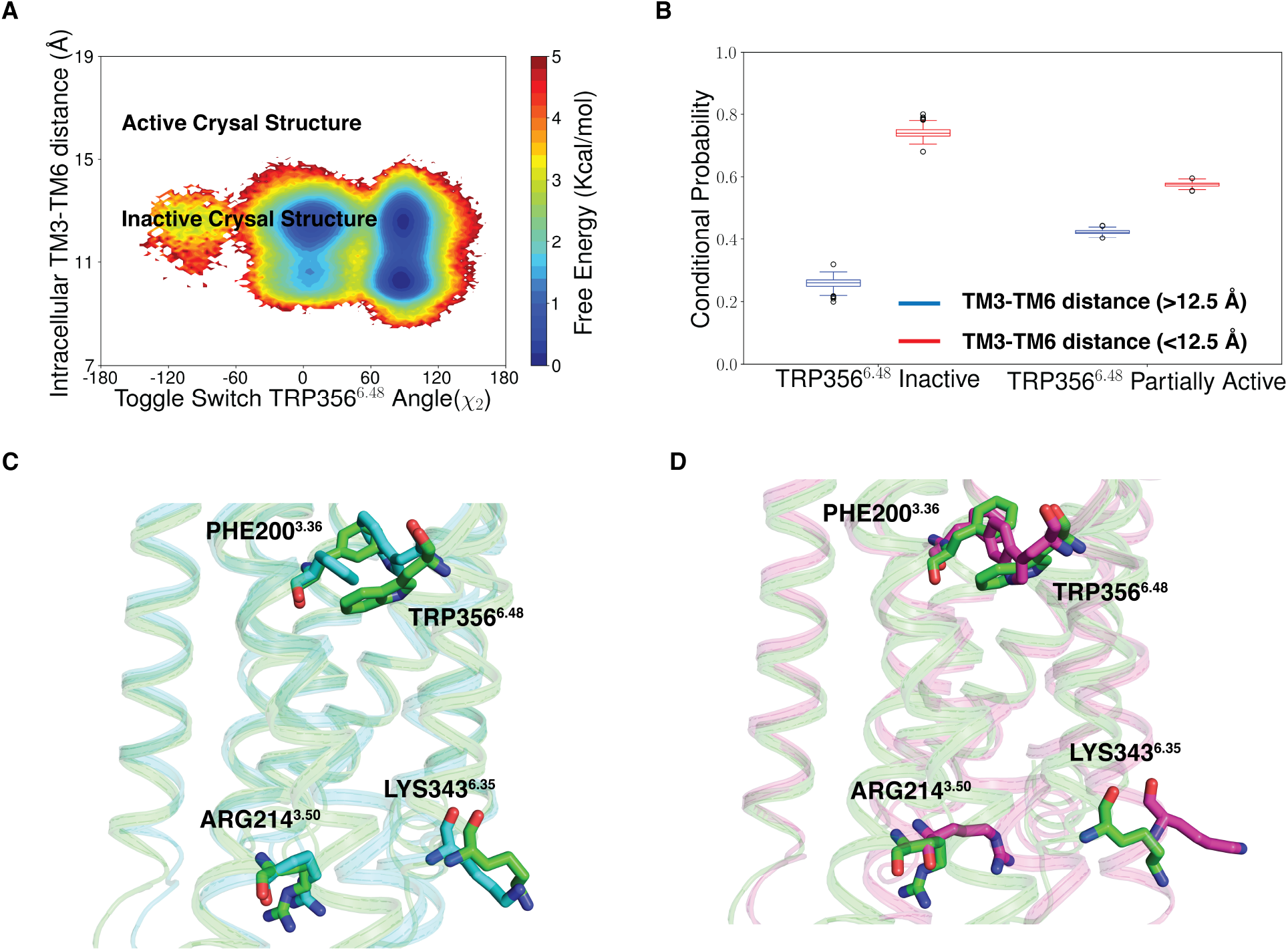
Movement of intracellular TM6 due to TRP356^6.48^ rotation (A) MSM weighted free energy landscape to capture toggle switch TRP356^6.48^ (TM6) *χ*_2_ angle and intracellular TM6 move-ment of CB_1_. Intracellular TM6 movement is measured ARG214^3.50^-C*α* (TM3) and LYS343^6.35^-C*α* (TM6). (B) Box plot to show the probabilities of intracellular TM6 movement with TRP356^6.48^ in inactive and partially active condition. Data distribution in box plot are generated with 200 rounds of bootstrap sampling with 80% of total number of trajectories (Method section). (C, D) Representative structure from partial active minima (color:green) is superimposed with inactive (color:cyan) (C) and active (color:pink) (D) structure of CB1 to depict the toggle switch movement and intracellular movement.

## Conclusion

In this study, we propose two hypotheses for the partial agonism of THC molecule for Cannabinoid receptor 1. Our first hypothesis is based on different binding position of the THC inside the orthosteric binding pocket. Second hypothesis states that THC may only able to move TM6 partially. To test the hypotheses, we perform unbiased molecular dynamics simulation for THC binding to CB_1_. Our results support both our hypotheses. Simulations show that during binding THC is stabilized in agonist-like and antagonist-like poses. While binding in the antagonist-like poses, the aromatic group of THC orients itself as Arm 2 of the antagonist, whereas alkylchain of THC can take the conformation of arm 1 and arm 3. During the binding process, alkyl chain orientation is shown to be important factor of determining the binding of THC to agonist pose. When THC enters the receptor with positive side chain dihedral angle (C2-C3-C1-C2), it leads to the upward movement of N-loop and favors the THC binding to the agonist-like pose. Whereas, with negative dihedral angle THC increases the free energy barrier for transition from antagonist-like binding pose to agonist-like binding pose. Therefore, chemical modification which can stabilize the side chain in perpendicular direction to the aromatic ring may increase the agonistic property of the ligand.

Over the years, various effects on THC alkyl chain modifications on binding and functionality have been proposed. Δ^9^-THC and Δ^8^-THC have similar binding affinity and agonistic property. It has been shown that Δ^8^-THC analog with cis double bond between C1 and C2 (which fixes the side chain in perpendicular direction) with same side chain length, increases the agonistic property of the ligand(*27*). Our simulations also support this experimental observation. With cis bond, THC analog can move to the agonist-like pose with less free energy barrier (Figure 4A) and therefore has more agonist property which bolsters our claim in the first hypothesis.

To generalize our first hypothesis that THC can be stabilized at both agonist and antagonist pocket, we performed docking of other partial agonists on representative CB1 structures pf THC bound agonist binding pose and antagonist binding pose (Figure S11). Five available partial agonists are selected for GPCRdb database(*29*): Magnolol, AM4089, NMP-4, (S)-Δ^3^-THC, (R)-Δ^3^-THC. Docking studies reveal stable docked poses for partial agonists in both the binding pocket. In the antagonist binding pose, we observed that the partial agonists extend downwards towards Na^+^ binding site similar to THC(*13*). These partial agonists binds with similar affinity in agonist and antagonist binding pocket. These docking results show that our hypothesis for partial agonism may be universally valid for other partial agonists for CB_1_.

Our results also show that the THC can rotate the toggle switch TRP356^6.48^ conformation when binding to the agonist like poses. The partially active conformation of CB_1_ matches well with CB2 inactive structure which helps to explain yin-yang relation for some ligands between CB_1_ and CB_2_. Recently published NMR-study on another class A GPCR, A_2_AAR, has shown that partial agonist rotates TRP246^6.48^ in a distinct conformation compared to full agonists or antagonists(*30*). These experimental observations support our results obtained computationally. Furthermore, our simulations reveal that partially active toggle switch movement consequently affects the intracellular side of the receptor and leads to the partial outward movement of TM6. In future study, this partial movement of the TM6 can be experimentally validated by DEER spectroscopy (*31-34*).

Overall, our present computational study with MD simulations and MSM is competent with previous experimental results and also provide new insights for partial agonism of THC for CB_1_. The findings of the paper are crucial for guiding the chemical modification for cannabinoid ligands for future drug development.

## Method

### System Preparation

Inactive structure of CB_1_ (PDB ID: 5TGZ(17)) is used as a starting structure for MD simulation. Non-protein residues and stabilizing fusion partner within the receptors third intracellular loop (ICL3) are removed from the PDB structure file. Hydrogen atoms are added to protein amino acid residues using reduce command of AMBER package (*35*). Truncated N-terminus and C-terminus & unconnected residues of TM5 and TM6 are capped with neutral terminal residues (acetyl and methylamide groups). Thermostabilized mutant residues are replaced with original residues using tleap (*17*). The modified protein structure is embedded in POPC bilayer using CHARMM-GUI (*36*). Salt concentration of 150 mM (Na+ and Cl-) is to neutralize the system. MD system is solvated using TIP3P water model. 3-D structure of THC is obtained from PubChem in sdf format. Forcefield parameters of THC are obtained using antechamber (*37, 38*). THC is added to MD system using packmol (*39*) to generate the starting structure.

### Simulation Details

Molecular dynamic simulations are performed using AMBER18. The MD system is subjected to minimization with gradient descent and conjugate gradient algorithm for 5000 and 10000 steps respectively. Minimized system is slowly heated from 0K to 10K and 10K to 300K in NVT ensemble to increase the temperature at desired level. Each step is done in 1 ns period. To control the pressure of the heated system at 1 bar NPT ensemble in employed. During modulation of temperature and pressure, protein backbone C*α* atoms are restrained with a spring force. Furthermore, total 50 ns of equilibration is performed in NPT ensemble to maintain the temperature and pressure at desired level of 300K at 1bar without any restraint force. Production runs are also performed in NPT ensemble. The Temperature and pressure of the system is maintained by berendsen thermostat and barostat (*40*). Simulation timestep of 2 fs is used for MD simulation. As, hydrogen atom is the lightest atom in the system, Vibrational frequency of the hydrogen can be lower than the specified timescale. It can create instability in the system due to the large fluctuation of hydrogen atom. Therefore, SHAKE algorithm(*41*) is used to put restraint in the movement of hydrogen atom by implementing lagrangian multiplier. For nonbonded force calculation, 10 Åcutoff distance is used. Periodic boundary condition is applied for all simulations. To consider the non-bonded long range interactions, Particle Mesh Ewald method is implemented(*42*). Simulations are performed with Amber FF14SB(*43*) and GAFF(*37*) forcefield parameters.

### Adaptive Sampling

Ligand binding to GPCR is rare event compared to MD simulations timescale. To capture the entire binding process, Adaptive sampling technique is utilized. This approach has been shown to sample the conformational ensemble of a variety of biological systems(*26, 44-46*) including the ligand binding process(*13, 47-50*). In this technique, data from one round of simulation is represented by conformational space of protein by using biological relevant features. The features are used to cluster the protein space using k-means clustering algorithm. The structures for next round of simulation are selected randomly from the cluster centers with lowest count of data points. This process is repeated in each round. For this case, intracellular and extracellular helical distance and THC binding distance from TM5 are used as adaptive sampling feature matrices. Although adaptive sampling helps to parallelize the simulation by sampling from lower probability space, it affects the ensemble distribution and therefore brings sampling error in free energy calculation. To overcome this caveat Markov state model (discussed below) is implemented to remove the sampling the bias from the simulation data. A total 143 *μ*s of simulation data was collected using adaptive sampling protocol.

### Markov State Model

The theory of MSM(*51-53*) depicts that in a sequence of event, the transition probability of moving from one state to another depend only on the present state and not on the path the system has taken to be there. The trajectory of Molecular dynamics simulation follows the same follows the same principle. According to the verlet algorithm, the evolution coordinate and momenta of atoms in system only depends on the on the present state. Therefore, MD trajectory can be assumed to be markovian and according to its nature, the probability distribution of the protein conformational ensemble can be calculated from the transition probability matrix between the states. Each element *T_ij_* of transition probability matrix calculates the probability of transition using the equation 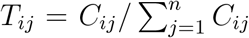 where *C_ij_* is the count of jump between the *i* and *j* and *C_i_* is the count of frame in state *i*. To make the transition probability matrix statistically significant, states with conformation are clustered together into microstates assuming there is no large energy barrier in the same microstate. However, clustering of conformational space increases the memory of each state which can invoke non-markovinity in our system. This issue can be overcome by increasing the lag time (*τ*) such that it preseves the markov property and hence satisfies the equation *p*(*t* + *τ*) = *p*(*t*)*T*(*τ*) where *p*(*t* + *τ*) and *p*(*t*) are vectors representing the probability of the microstates. Pyemma(*54*) package is used to construct MSM. We select 23 biologically important features (Table S1) to capture the THC binding and protein conformational changes. Features are transformed into time-lagged independent components (tics) (*55, 56*) to find the slowest components. Tic components are shown to correlate well with important features for THC binding (Figures S12A, S12B, and S12C). The transformed data is clustered using k-means clustering algorithm. For MSM building, lag time is chosen by finding out logarithmic convergence of process timescales computed from MSM eigenvalues (process timescale 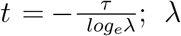 is the eigenvalue) (Figure S13). To find out the optimal number of clusters and optimal tic variance, MSM is subjected to VAMP2(*55*) scoring to measure the kinetic variance (Figure S14). Based on Transition matrix, MSM predicts the population (stationary probablity, *π_i_*) of each clustered state which is needed calculate thermodynamics (*G_i_* = – *k_b_T log_e_π_i_*) of the process.

### Transition path theory

To obtain the kinetics for transition between MSM macrostates, transition path theory (*57, 58*) is implemented. TPT calculates the flux between two macrostates (which consists of one or multiply MSM states) using MSM transition matrix (*59*). The Flux (*FAB*) is given by the equation 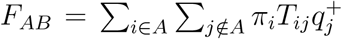 where 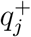 is probability that state j will reach macrostate B before A (*60*). From the flux calculation, we can compute mean free passage time (MFPT) using equation 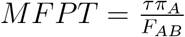. where *π_A_* is the probability the system was in macrostate A. TPT calculation is performed with Pyemma (*54*) package. Here, the macrostates are defined manually. Five MSM states with highest raw count inside area of interest, is defined as a macrostate. For example, macrostate 4 in Figure 5A is consisted of five MSM states with highest raw count inside the area where THC(C1’)-TYR275^5.39^(C*α*) distance is between 9 *A* and 13 ^Å^ & THC didehral angle is between 60° and 120°.

### Trajectory analysis

All the feature calculation and data processing from the MD simulation was done CPPtraj(*61*) and MDtraj (*62*). For the trajectory visualization and analysis, VMD (*63*) and pymol v1.7 (Schrodinger, LLC) software package was used. To calculate ligand and protein residue contact, GetContacts package (https://getcontacts.github.io/) was implemented.

### Kinetic Monte Carlo simulation

Kinetic monte carlo (kMC) simulation is a stochastic method to visualize the evolution of a system based on probability of the transition between different states. In this case, transition probability of different states was obtained from MSM. There are few steps to implement kMC in MSM weighted MD data. First, we generate a random number (R) between 0 to 1. Second, we describe the possible transition event from i as discrete cumulation probability distribution. Therefore, the cumulative transition probability between *i* to *j* can be written as 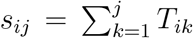. If R lies between *s_i,j_* and *s*_*i,j*+1_ then system transition happens from *i*th to *j*th state. Time required for each transition is taken to be the lag time considered for MSM. Same procedure is repeated desired number of steps. To build a trajectory from kMC state evolution, we pick a random frame at each step from the chosen state. As we are interested to observe the ligand binding dynamics, kMC simulation is started from a state where THC was 25 Å away from TYR275^5.39^ (C*α*).

### Docking Study

Docking study was performed using Auto dock vina (*64*) software. 3-D structures of partial agonists are selected from PubChem in sdf format. Antechamber is used to convert the ligands in mol2 format and add partial changes. Then, Auto dock is used to convert the ligands into required pdbqt format. For the docking of the ligands in both agonist and antagonist pose, the box size is calculated from THC binding structure in respective poses. To dock the ligand in antagonist and agonist bound poses, MD predicted structures are used.

### Error Analysis

To determine error in our thermodynamics (Free energy, Conditional probability) and kinetics (TPT) calculations we perform bootstrap analysis on MD data (*65*). In each bootstrap sample, we randomly pick N trajectories, where N is equal to 80% of total number of trajectories. We keep the state labeling same as the original MSM. For every bootstrap sample, MSM is computed to determine thermodynamics and kinetics. Total 200 bootstrap samples are generated for error calculations.

