## Supplementary Information for "Why does Δ^9^-Tetrahydrocannabinol act as a partial agonist of cannabinoid receptors?"

### Supporting information: Why $\triangle^9$-Tetrahydrocannabinol acts as a partial agonist of cannabinoid receptors?

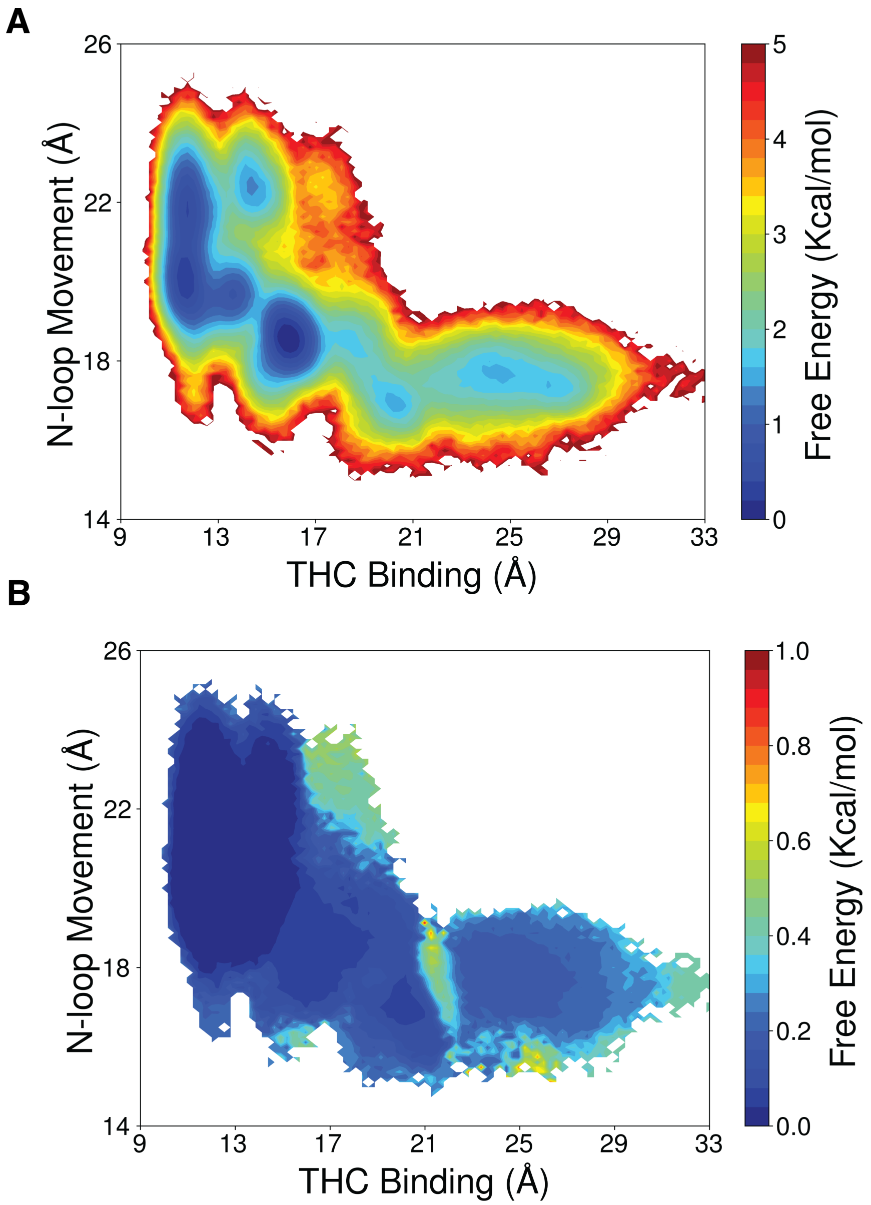


Figure S1: (A) Unweighted free energy landscape to capture THC binding and N-loop upward motion. (B) Errors on the free energy landscape projected on the same matrices. THC binding distance is measured between THC-C1' and TYR275$^{5.39}$-C$\alpha$ (TM5) and N-loop upward motion is measured between MET103$^{\mathrm{N-loop}}$-C$\alpha$ (N-loop) and ASP163$^{2.50}$-C$\alpha$ (TM2). Errors are calculated with 500 rounds of bootstrap sampling with 80% of total number of trajectories (Method section).

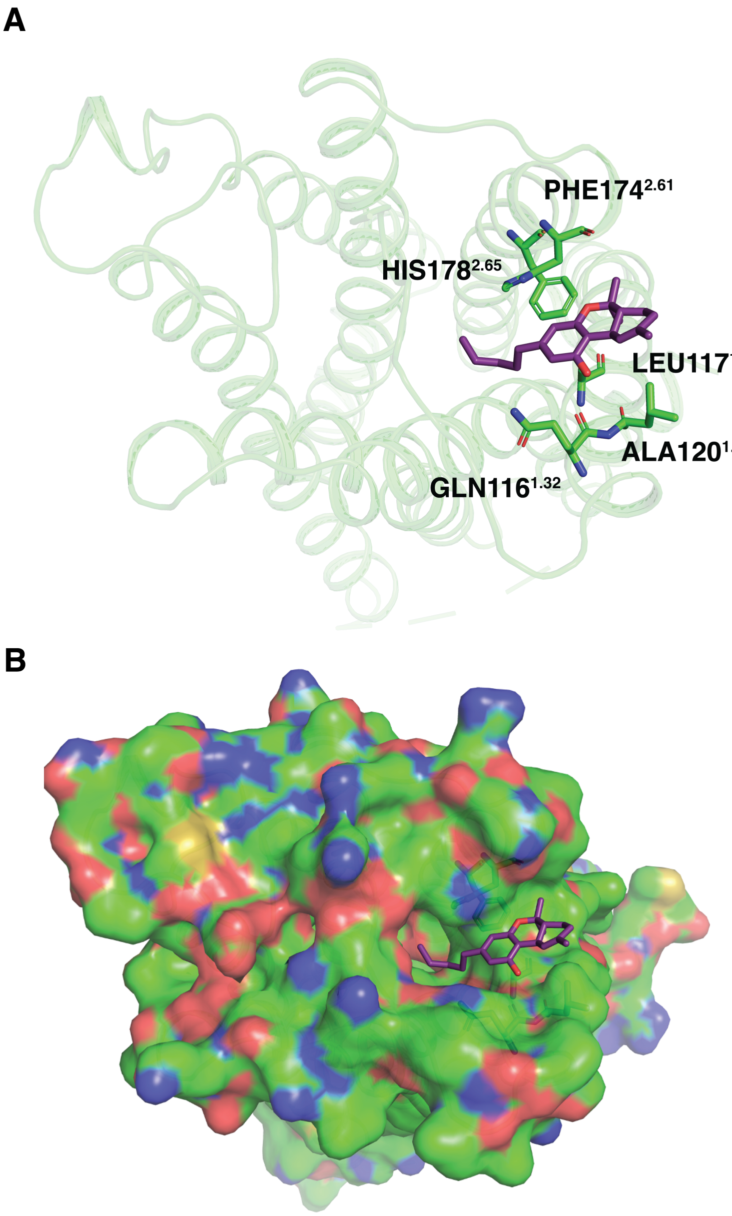


Figure S2: MD snapshot representing binding of THC from the extracellular region through the space between N-loop, TM1 and TM2 (top view). Proteins are represented as cartoon in (A) and as surface in (B). THC (color:violet) and interacting residues (color: green) are shown as stick representation.

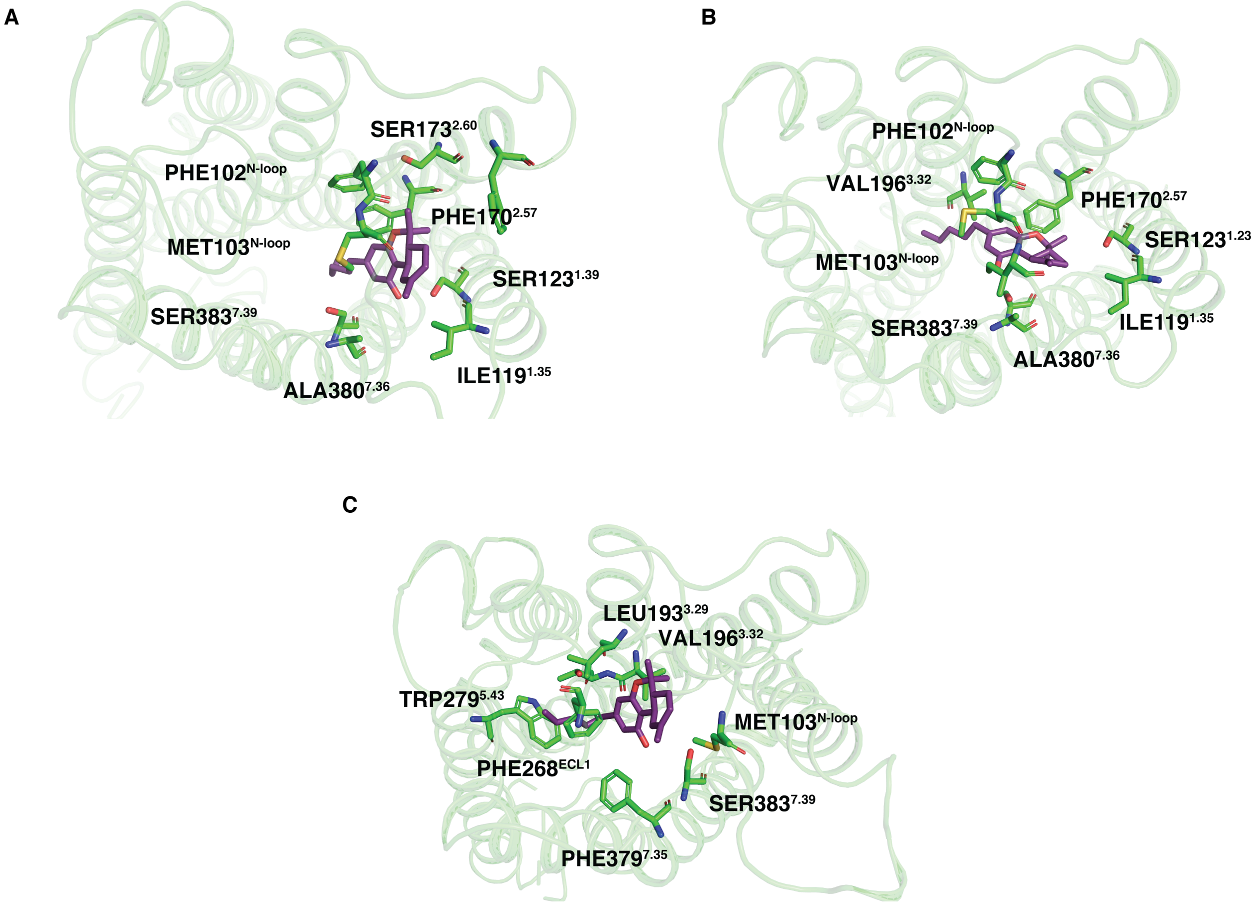


Figure S3: Important interactions between protein residues and THC at different stabilized positions during binding (top view). (A), (B) shows a representative structure from antagonist-like pose 1 and 2, respectively. (C) shows a representative structure from agonist-like pose. Stable interactions were measured using GetContacts package. Protein structures are shown as cartoon representation (color: green). THC (color: violet) and interactive residues (color: green) are shown as stick.

A

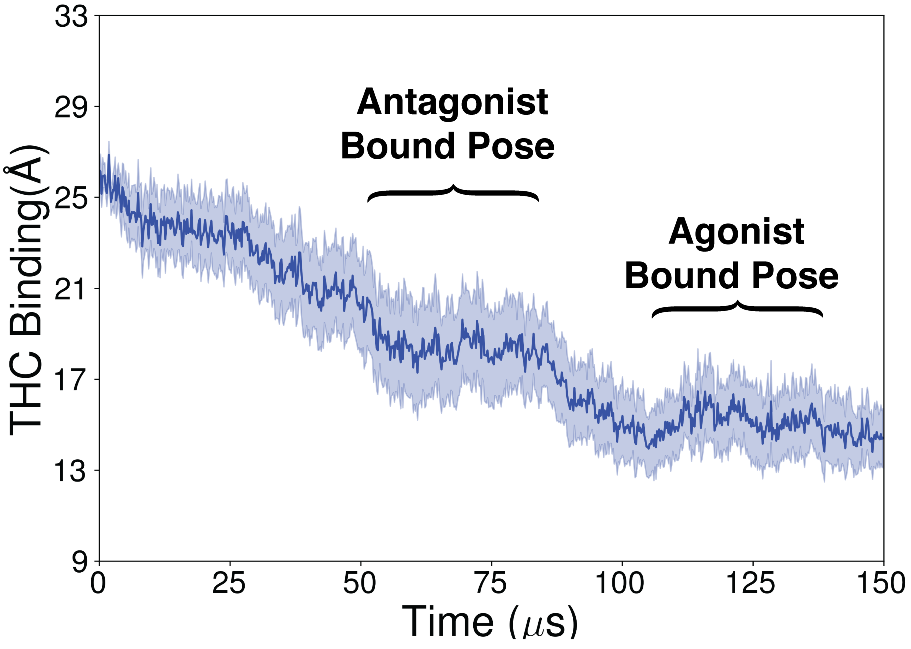


B

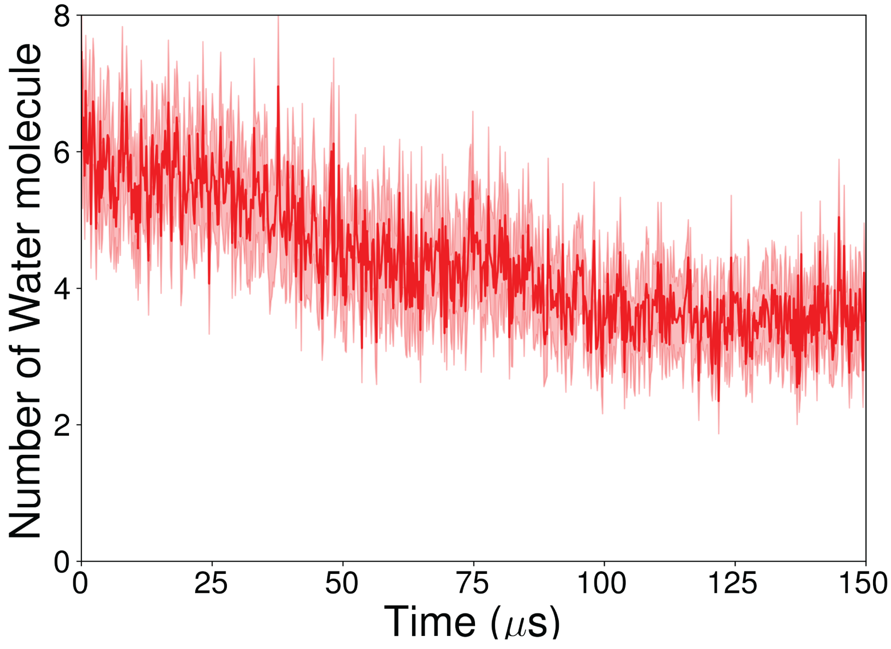


Figure S4: Kinetic Monte Carlo trajectory of 150 $\mu$s generated with the THC outside of the receptor. THC-C1' and TYR275$^{5.39}$-C$\alpha$ (TM5) distance (A) and number of water molecules surrounding the the ligand (B) plotted against the time to show the transition during binding. Error bars are shown as semitransparent region. Error bars in the plot are calculated by running kMC simulations for 20 times.

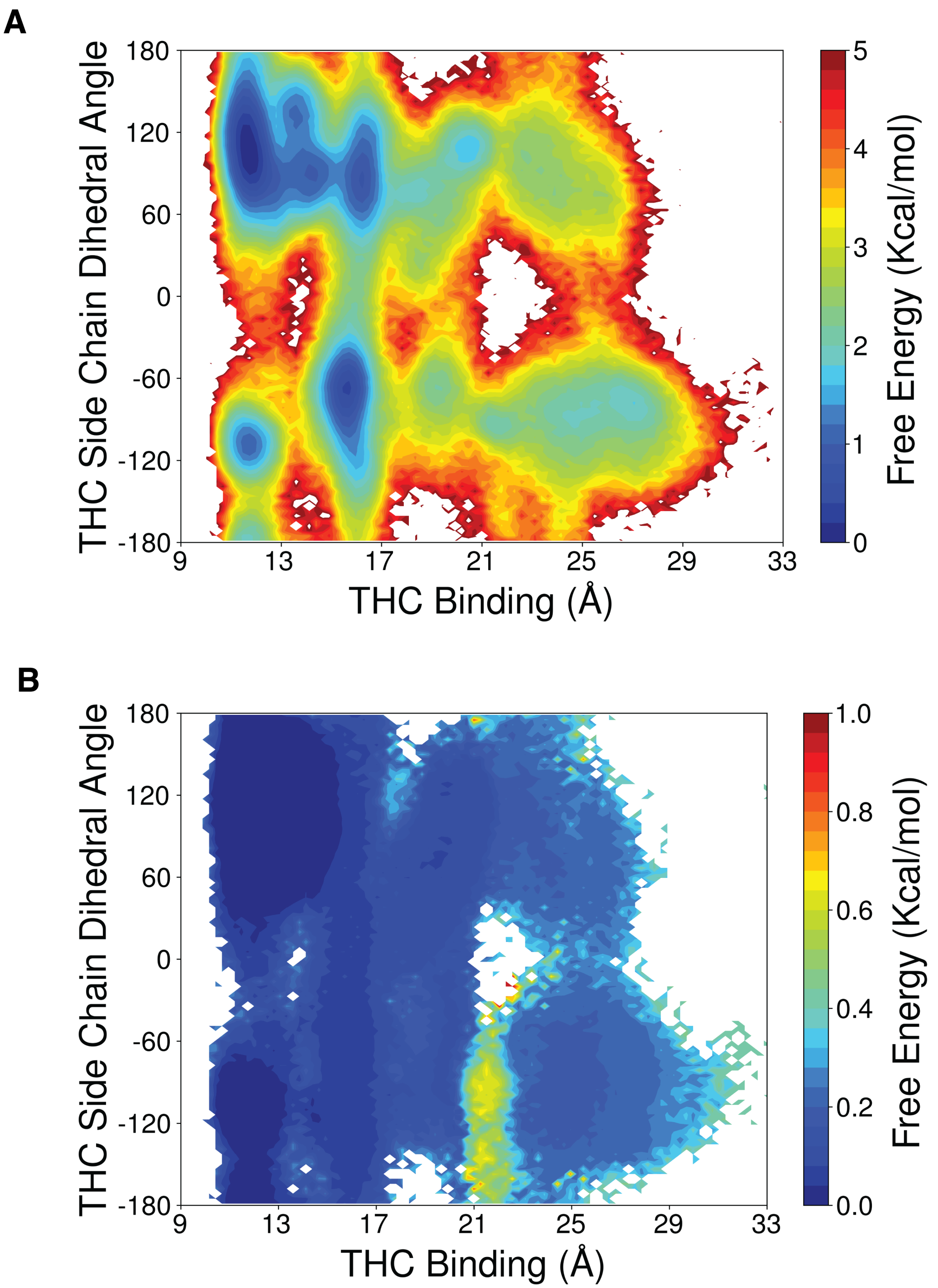


Figure S5: (A) Unweighted free energy landscape to capture THC binding and THC side chain dihedral. (B) Errors on the free energy landscape projected on the same matrices. THC binding distance is measured between THC-C1' and TYR275$^{5.39}$-C$\alpha$ (TM5) and THC sidechain dihedral is measured between C2,C3,C1',C2'. Errors are calculated with 500 rounds of bootstrap sampling with 80% of total number of trajectories (Method section).

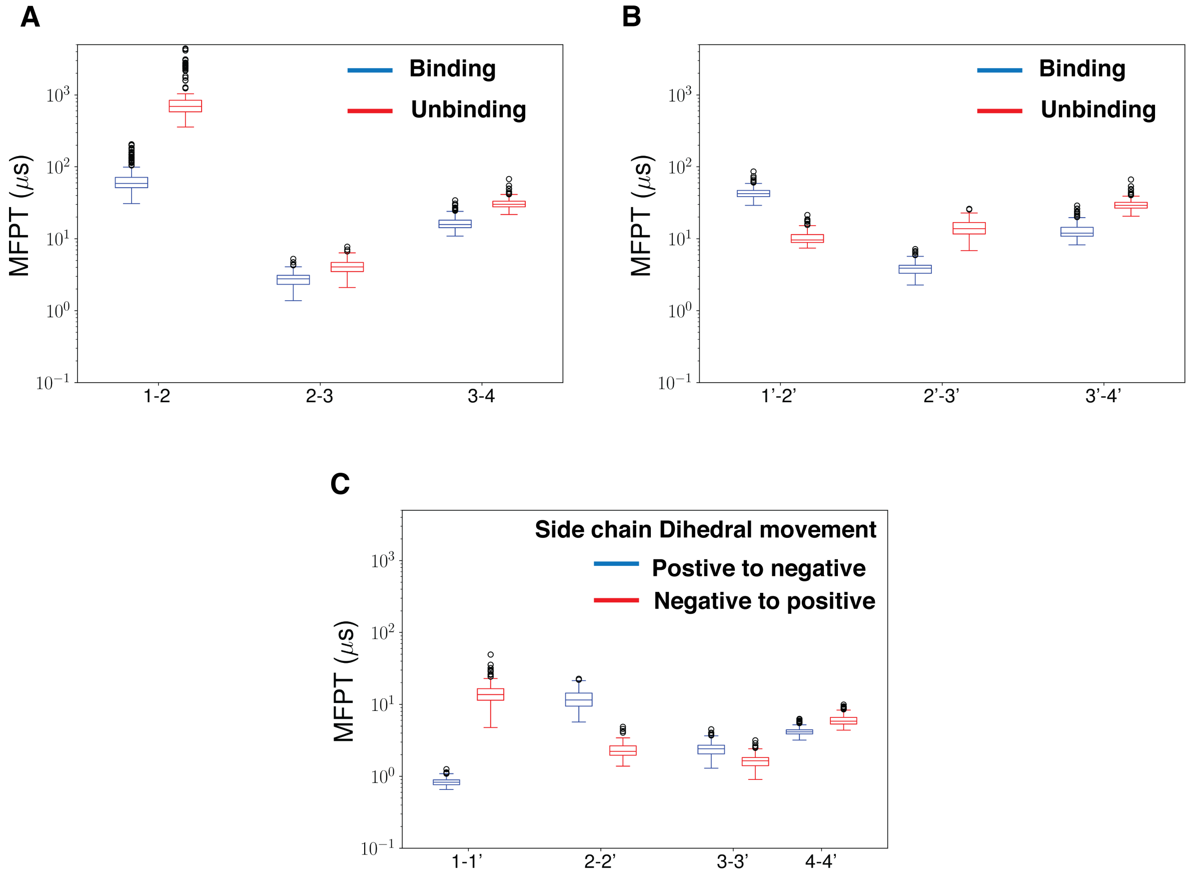


Figure S6: Box plot for mean free passage time between the macrostates as shown in Figure 4A. (A), (B) are showing the transitions between macrostates in top and bottom panel, respectively. Blue boxes are representing binding and red boxes are representing unbinding. (C) is representing MFPTs for dihedral transitions between macrostates with positive and negative dihedral.

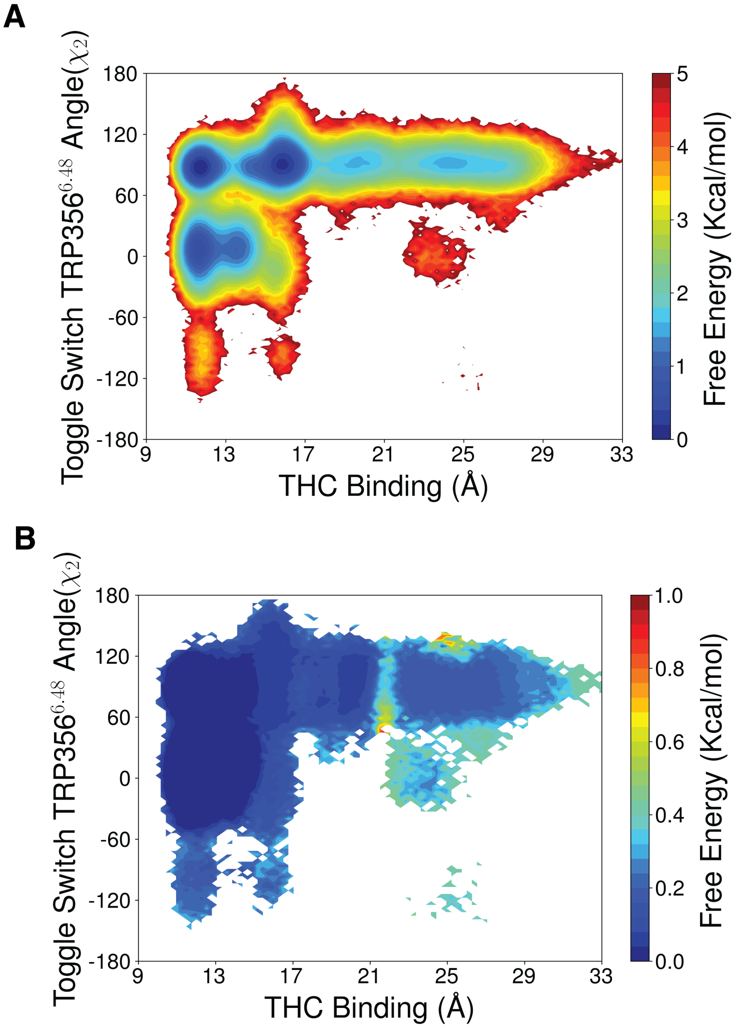


Figure S7: (A) Unweighted free energy landscape to capture THC binding and toggle switch TRP356 (helix 6) $\chi_2$ angle. (B) Errors on the free energy landscape projected on the same matrices. THC binding distance is measured between THC-C1' and TYR275$^{5.39}$-C$\alpha$ (TM5). Errors are calculated with 500 rounds of bootstrap sampling with 80% of total number of trajectories (Method section).

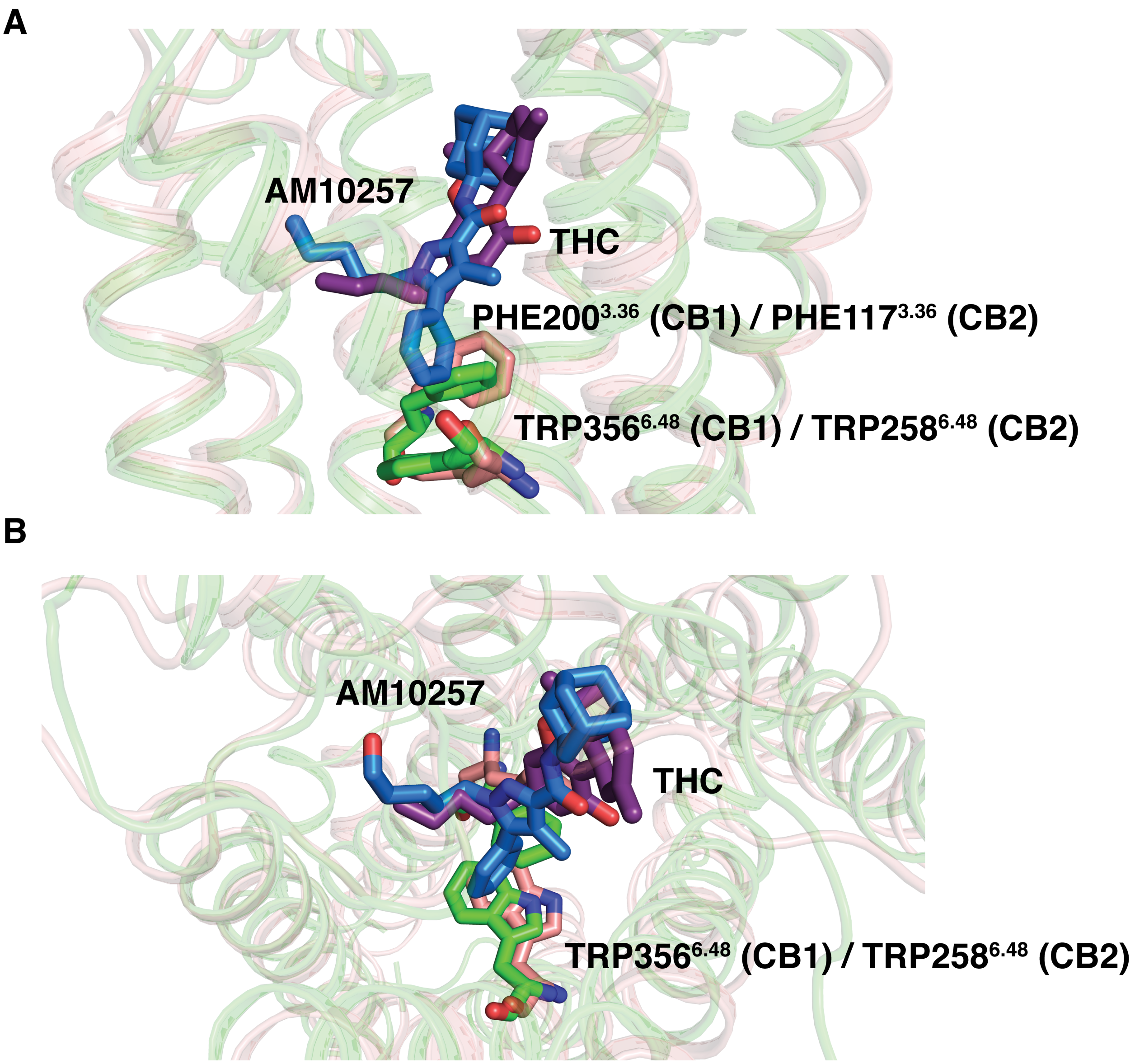


Figure S8: Comparison of partially active (state 1) (color: green) structure with inactive $CB_2$ (color: light brown) from side (A) and top view (B). Toggle switch residues and ligands (AM10257: blue,THC: violet) are shown as stick representation.

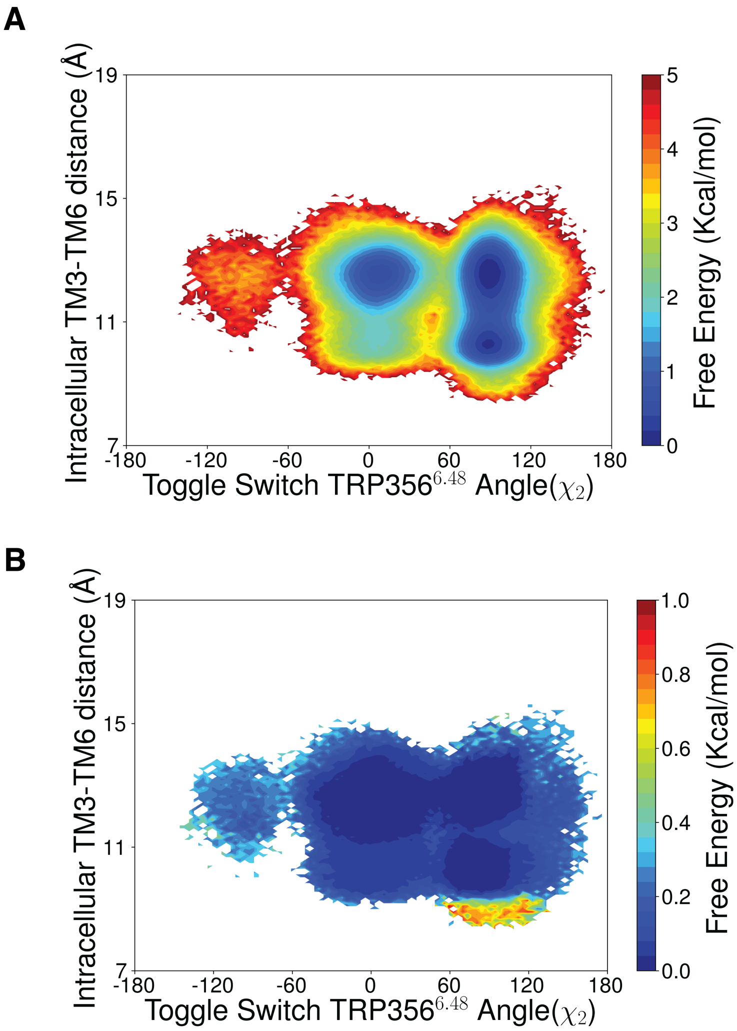


Figure S9: (A) Unweighted free energy landscape to capture toggle switch TRP356 (helix 6) $\chi_2$ angle and intracellular TM6 movement of CB1. (B) Errors on the free energy landscape projected on the same matrices. Intracellular TM6 movement is measured ARG214$^{3.50}$-C$\alpha$ (TM3) and LYS343$^{6.35}$-C$\alpha$ (TM6). Errors are calculated with 500 rounds of bootstrap sampling with 80% of total number of trajectories (Method section).

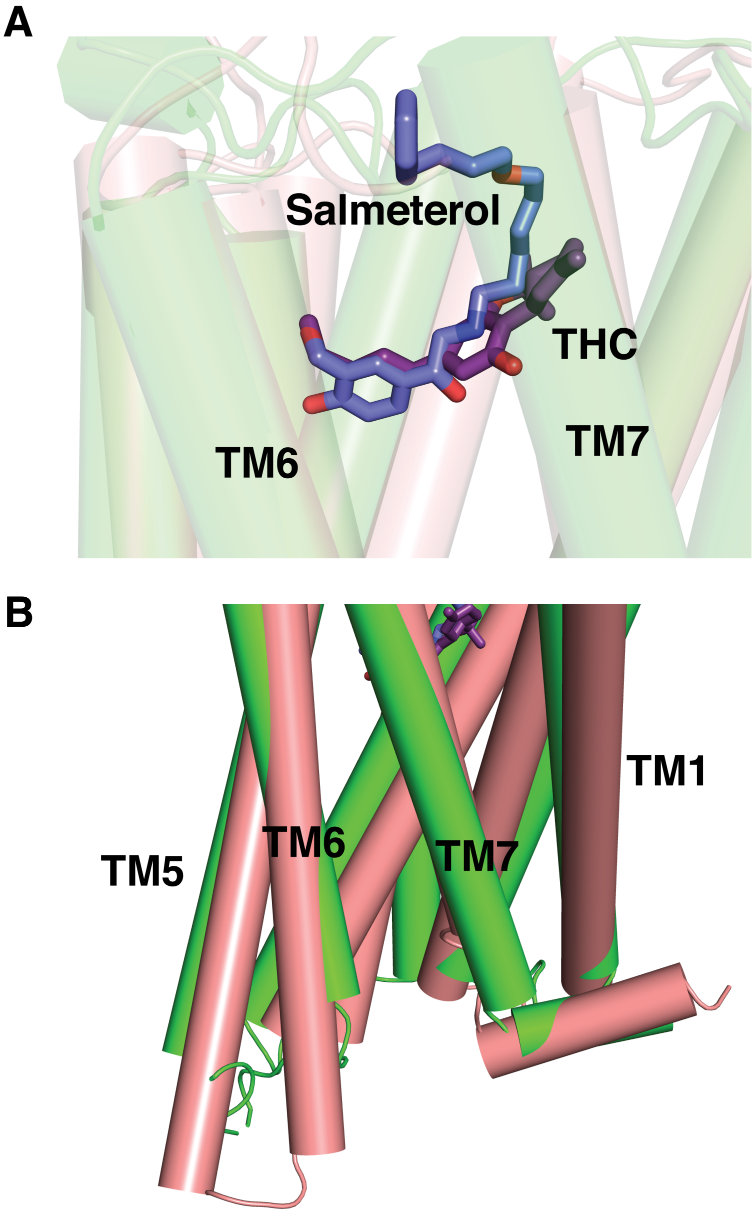


Figure S10: Comparison of partially active $CB_1$ (state 1) (color: green) structure with partially active $\beta_2$-AR (PDB ID: 6CSY, color: light brown) from side view. (A) focuses on the comparison of ligand bound pose of THC (color: violet) and Salmeterol (color: blue). (B) shows the TM orientation of both the receptor. ligands (salmeterol: blue, THC: violet) are shown as sticks.

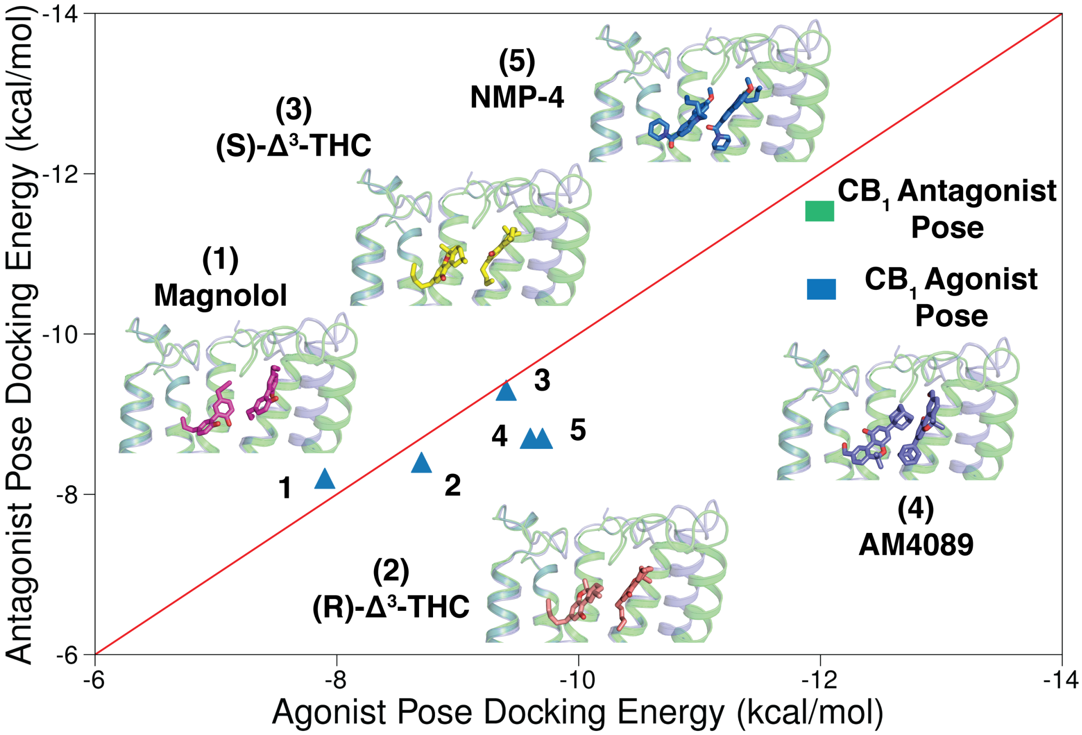


Figure S11: Agonist pose docking energy is plotted against antagonist pose docking energy for five different partial agonists. Protein with docked partial agonists in antagonist pocket(color: green) and agonist pocket (color: purple) are superimposed on each other. Five partial agonists are represented as stick with different color (Magnolol:pink,(R)-$\Delta^3$-THC: light brown,(S)-$\Delta^3$-THC: yellow, AM4089: purple, NMP-4: blue).

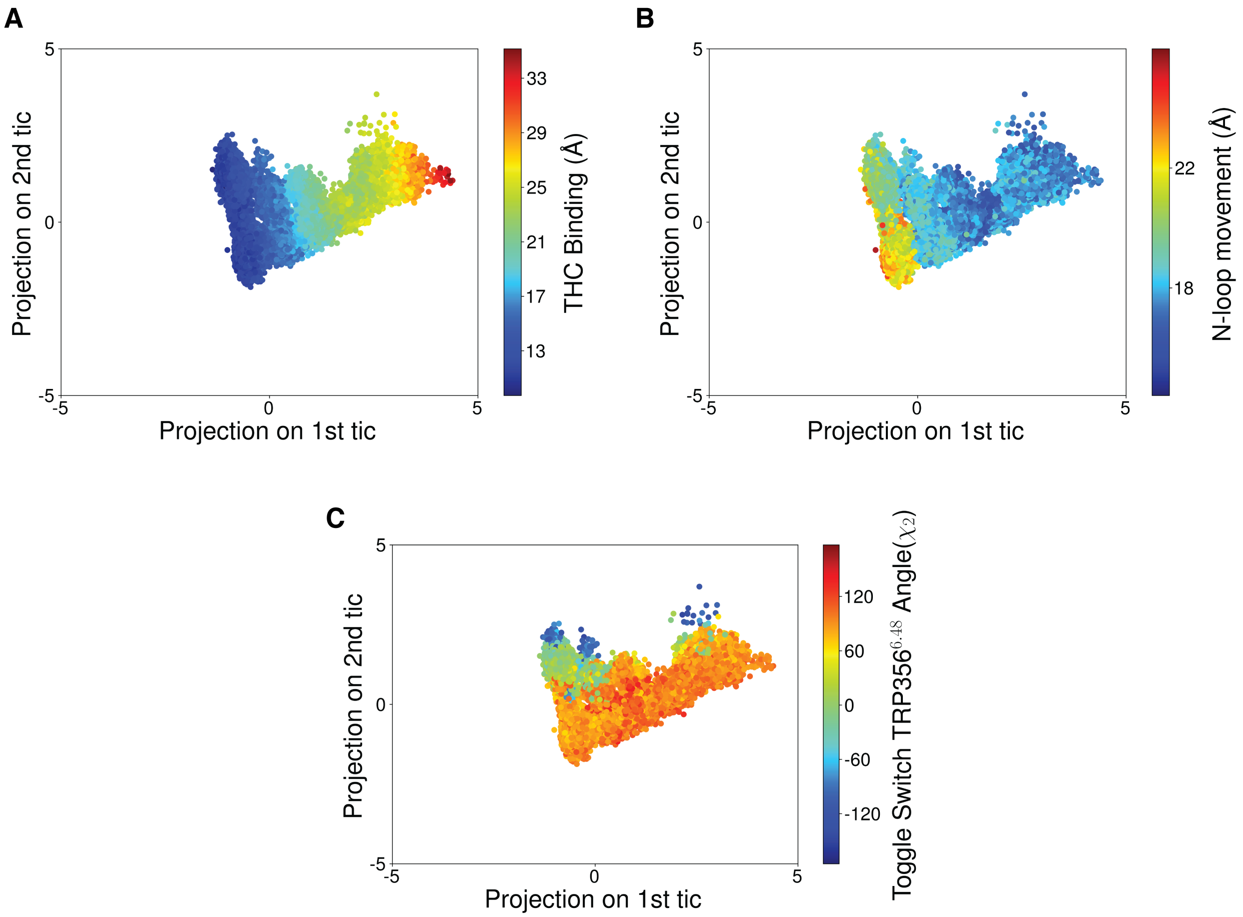


Figure S12: Correlation of 1st and 2nd tic components with system features. First tic component is highly correlated with THC binding (A) and N-loop movement (B). Second tic component is highly correlated with TRP356$^{6.48}$ $\chi_2$

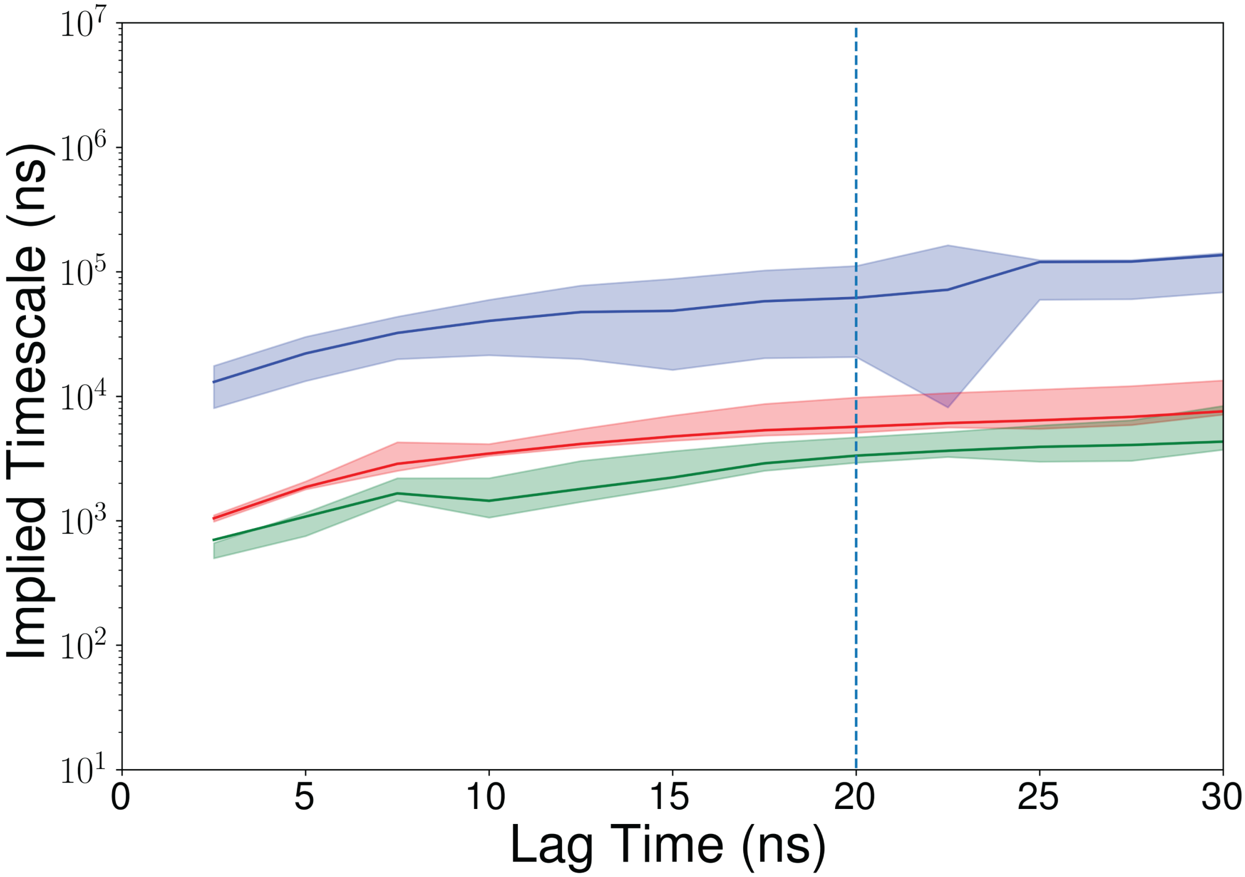


Figure S13: Logarithmic convergence of timescales ($2^{nd}$, $3^{rd}$ and $4^{th}$ highest eigenvalues) plotted against various lag times. Lag time of 20 ns used to build our MSM. Errors are calculated with 10 rounds of bootstrap sampling with 80% of total number of trajectories (Method section).

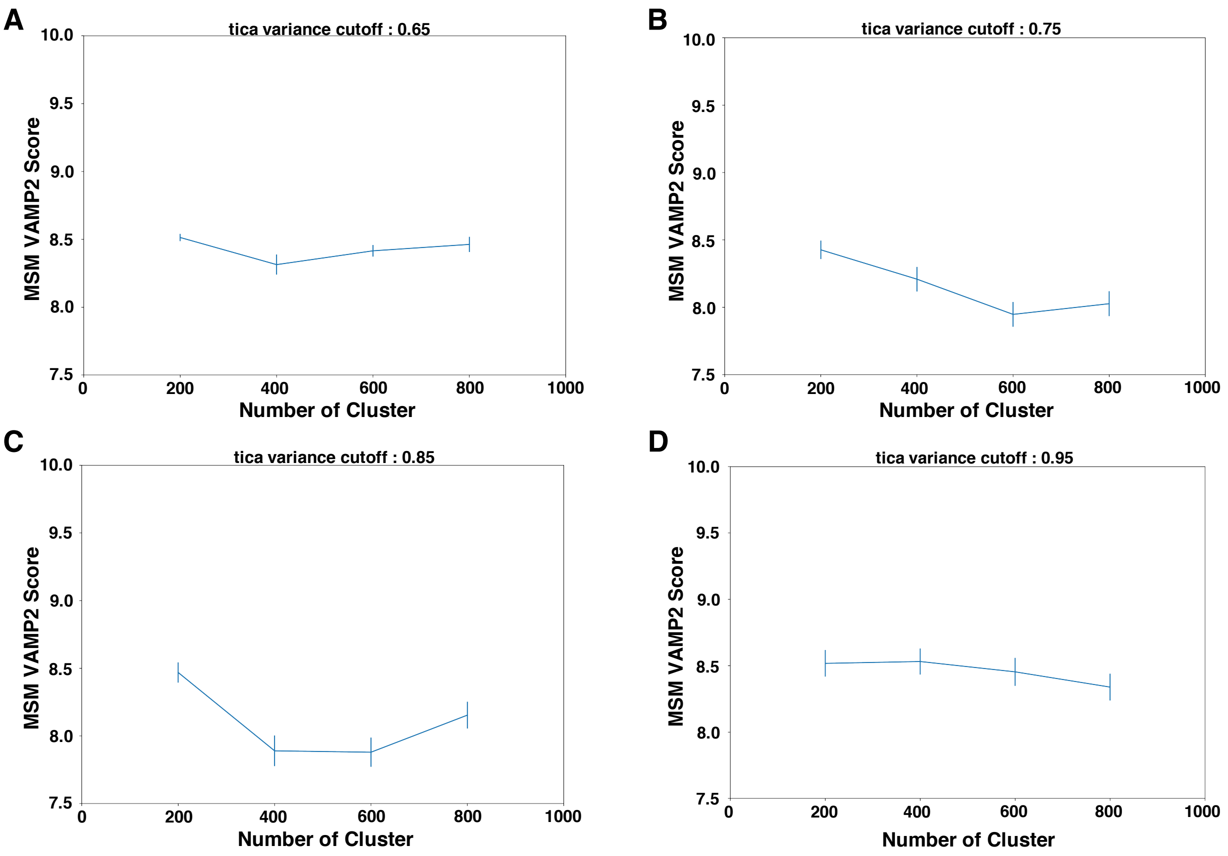


Figure S14: Optimal cluster number are chosen based on VAMP2 score of MSM based on different tic variance cutoff. MSM with tic variance of 95% and 200 clusters were selected for our simulation.

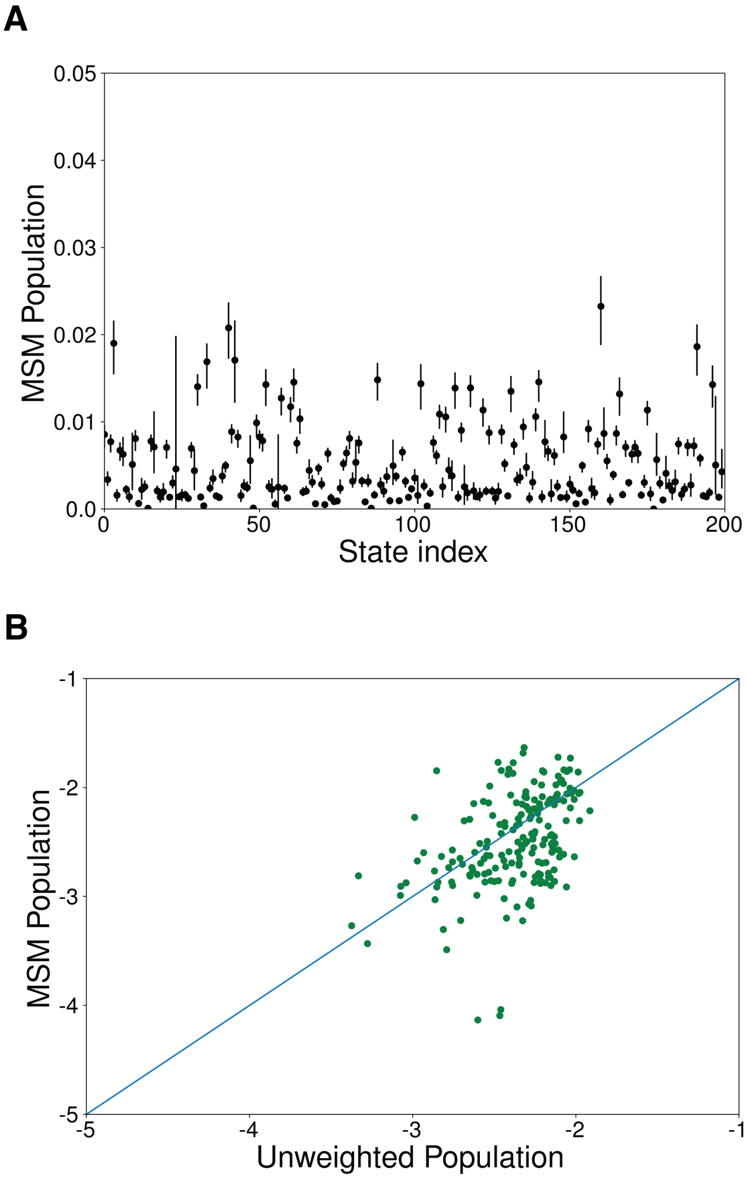


Figure S15: (A) Probability density of each MSM state. (B) MSM population vs Raw count in each clustered state. Errors are calculated with 10 rounds of bootstrap sampling with 80% of total number of trajectories (Method section).

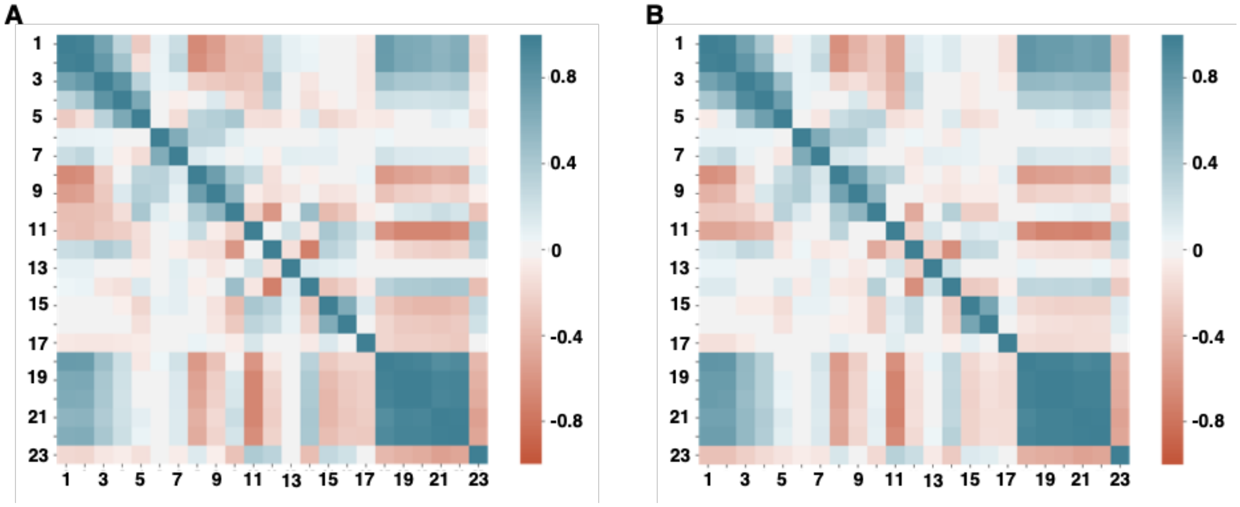


Figure S16: (A) Unweighted and (B) MSM weighted co-relation matrix of the features used to build MSM. Features corresponding feature number can be found from Table 1.

Table S1: Distance and angle features used to build Markov state model.

| Position | Number | Type | Feature |
|---|---|---|---|
| Extracellular helical movement | 1 | distance | $ASN116^{TM1}$(CA)-$PHE189^{TM3}$(CA) |
| | 2 | distance | $ASN116^{TM1}$(CA)-$PRO251^{TM4}$(CA) |
| | 3 | distance | $ASN116^{TM1}$(CA)-$LEU276^{TM5}$(CA) |
| | 4 | distance | $ASN116^{TM1}$(CA)-$ASP366^{TM6}$(CA) |
| | 5 | distance | $ASN116^{TM1}$(CA)-$THR377^{TM7}$(CA) |
| | 6 | distance | $ASP176^{TM2}$(CA)-$PHE189^{TM3}$(CA) |
| | 7 | distance | $ASP176^{TM2}$(CA)-$PRO251^{TM4}$(CA) |
| | 8 | distance | $ASP176^{TM2}$(CA)-$LEU276^{TM5}$(CA) |
| | 9 | distance | $ASP176^{TM2}$(CA)-$ASP366^{TM6}$(CA) |
| | 10 | distance | $ASP176^{TM2}$(CA)-$THR377^{TM7}$(CA) |
| N-loop movement | 11 | distance | $MET103^{N-loop}$(CA)-$ASP163^{TM2}$(CA) |
| | 12 | distance | $MET103^{N-loop}$(CA)-$PHE268^{ECL2}$(CA) |
| Toggle switch movement | 13 | Dihedral Angle ($\chi_2$) | $PHE200^{TM2}$ |
| | 14 | Dihedral Angle ($\chi_2$) | $TRP356^{TM6}$ |
| Intracellular helical movement | 15 | distance | $PHE155^{TM2}$-$LYS343^{TM6}$ |
| | 16 | distance | $ARG214^{TM3}$-$LYS343^{TM6}$ |
| | 17 | distance | $TYR153^{TM2}$-$TYR397^{TM7}$ |
| THC binding | 18 | distance | THC(C3)-$PHE268^{ECL2}$ |
| | 19 | distance | THC(C1')-$TYR275^{TM5}$ |
| | 20 | distance | THC(C1')-$TRP279^{TM5}$ |
| | 21 | distance | THC(C5')-$TYR275^{TM5}$ |
| | 22 | distance | THC(C5')-$TRP279^{TM5}$ |
| | 23 | Dihedral Angle (C2-C3-C1'-C2') | THC |

Table S2: Amount of simulation ($\mu$s) in each round of adaptive sampling

| Sampling Round | Total time |
|---|---|
| 1 | 27.83 |
| 2 | 4.83 |
| 3 | 4.68 |
| 4 | 9.18 |
| 5 | 5.78 |
| 6 | 4.13 |
| 7 | 11.93 |
| 8 | 11.49 |
| 9 | 11.44 |
| 10 | 5.13 |
| 11 | 4.93 |
| 12 | 4.73 |
| 13 | 2.11 |
| 14 | 6.68 |
| 15 | 5.37 |
| 16 | 2.57 |
| 17 | 2.73 |
| 18 | 7.98 |
| 19 | 9.49 |
| Total time | 143.01 |
